# Connectivity networks and delineation of distinct coastal provinces along the Indian coastline using large-scale Lagrangian transport simulations

**DOI:** 10.1101/2021.04.24.441108

**Authors:** D. K. Bharti, Katell Guizien, M. T. Aswathi-Das, P. N. Vinayachandran, Kartik Shanker

## Abstract

Ocean circulation defines the scale of population connectivity in marine ecosystems, and is essential for conservation planning. We performed Lagrangian transport simulations and built connectivity networks to understand the patterns of oceanographic connectivity along the Indian coastline. In these networks, nodes are coastal polygons and the edges connecting them represent the magnitude of larval transfer between them. We assessed the variation in connectivity networks within and between two monsoonal seasons, across El Niño–Southern Oscillation (ENSO) years and for pelagic larval durations (PLD) up to 50 days. We detected well-connected communities, mapped frequent connectivity breaks and ranked coastal areas by their functional role using network centrality measures. Network characteristics did not differ based on the ENSO year, but varied based on season and PLD. Large scale connectance (entire Indian coastline) was small, ranging from 0.5% to 3.4%, and the number of cohesive coastal communities decreased from 60 (PLD <4 days) to 30 (PLD >20 days) with increasing PLD. Despite intra-seasonal variation in connectivity breaks, four disconnected provinces were consistently identified across the entire PLD range, which partially overlapped with observed genetic and biogeographic breaks along the Indian coastline. Our results support the adoption of an adaptive regional management framework guided by fine-scale analysis of connectivity within the four provinces delineated in the present study. A few sites within each province displayed notably higher centrality values than other nodes of the network, but showed variation with season and PLD, and could be targeted for national and transnational conservation and management plans.

## Introduction

Marine populations have traditionally been described as ‘open’, with high exchange of propagules between spatially distant populations owing to oceanic transport of pelagic life stages (Roughgarden et al., 1988). However, it is becoming apparent that local oceanographic features and species-specific traits determine where a population falls in the continuum from ‘open’ to ‘closed’ (Levin, 2006). Apart from influencing demographic processes at ecological time-scales (Gaines et al., 2007), long distance transport using ocean currents can also have consequences over evolutionary time-scales (Jablonski, 1986; Paulay & Meyer, 2006). This would shape patterns of population genetic connectivity across taxonomic groups and large-scale patterns of diversity and community structure.

An understanding of oceanographic processes is therefore important in the study of marine dispersal. However, field studies of pelagic larval transport are prohibitively difficult because of small larval size, challenges in taxonomic identification and navigating the vastness of the open ocean (Pineda et al., 2007). Biophysical modelling combines ocean circulation simulations with larval traits and habitat information to provide estimates of larval transport. The latter can be compared to empirical patterns of gene flow between populations to understand the demographic effects of larval transport on marine population connectivity (Liggins et al., 2013).

The accuracy of larval transport estimates depend upon the resolution of ocean circulation simulations (Briton et al. 2018), location of reproductive populations and the degree of species-specificity in larval release timing, pelagic larval duration (PLD) (Di Franco & Guidetti, 2011), and larval behaviour (Guizien et al. 2006). Although larval behaviour can strongly alter dispersal patterns and distribution of suitable habitat can restrict connectivity, ocean flow and PLD are the minimum parameters required to assess seascape connectivity (North et al., 2009). Such general biophysical simulations of larval transport based on global ocean circulation simulations can serve as a foundation for regional ecological studies by delineating disruptions in connectivity (Rossi et al., 2014). Within well connected regions, the frequent exchange of individuals among spatially distinct populations can influence their demography, giving rise to a metapopulation (Hanski & Gagiotti, 2004). Identifying the extent of well-connected populations, within which metapopulation functioning is expected to occur, is a pre-requisite for establishing the scale of biodiversity management (Halpern & Warner, 2003). Since management policies are a national prerogative, it is also important to include all the regions falling within the boundaries of an Exclusive Economic Zone (EEZ) in connectivity studies.

The EEZ of India includes (1) a coastline of over 5000 km that divides the northern Indian Ocean into two basins – the Arabian Sea in the west and the Bay of Bengal in the east, (2) the Lakshadweep archipelago located in the Arabian Sea, and (3) the Andaman and Nicobar archipelago located in the Bay of Bengal. Sri Lanka is an island located in the Bay of Bengal, which is separated from India by the narrow Palk Strait, where both countries have contiguous EEZs (Figure 1). There is an absence of large-scale connectivity studies from the northern Indian Ocean (see George et al., 2011 and Gaonkar et al., 2012 for local-scale studies) despite a large body of physical oceanography studies from this region (reviewed in Shetye & Gouveia, 1998; Schott & McCreary, 2001; Shankar et al., 2002; Schott et al., 2009; Vinayachandran, 2009; Hood et al., 2017).

**Figure 1.**
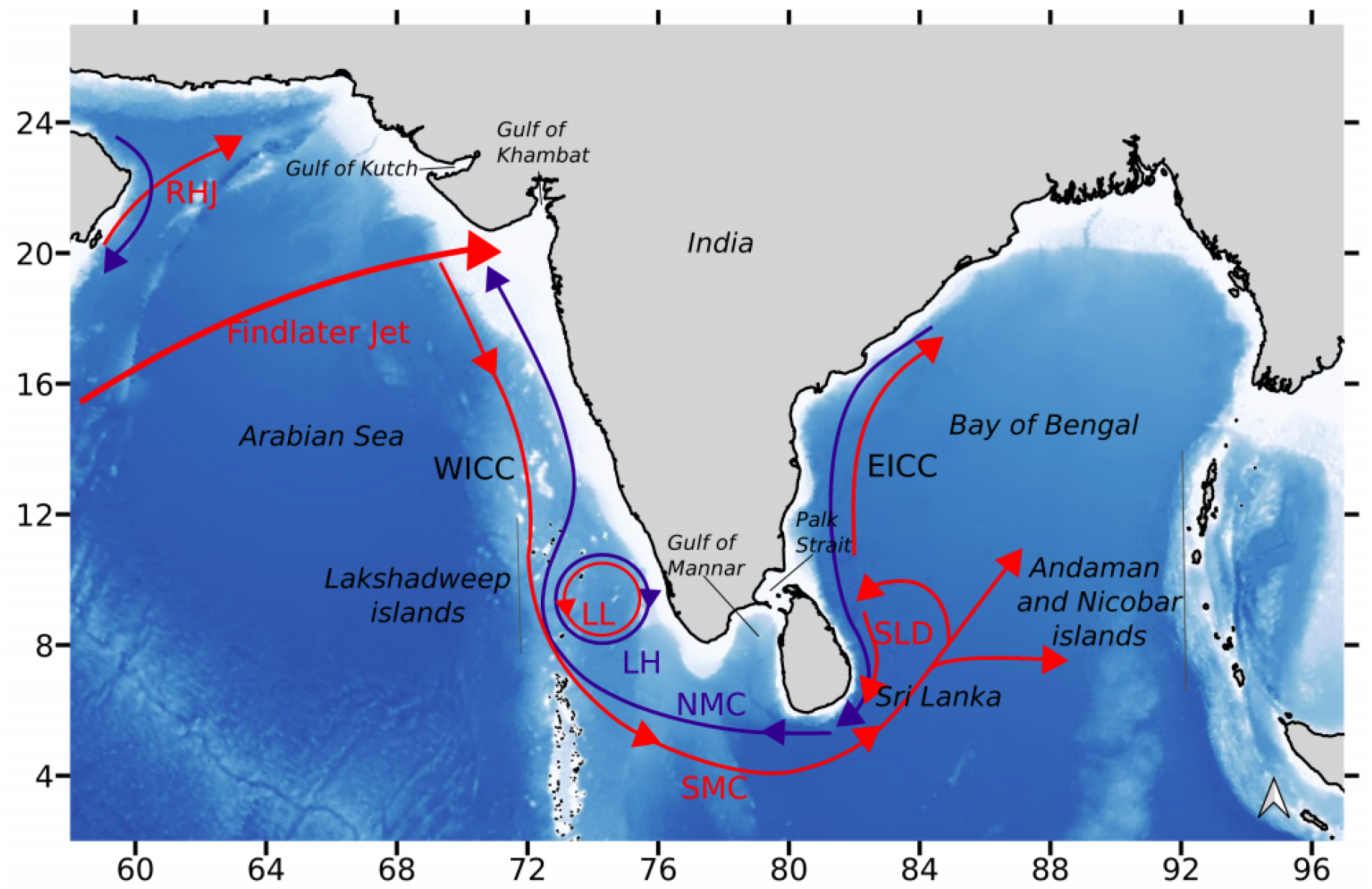
Schematic illustration of major currents along the Indian coastline – Ras al Hadd Jet (RHJ), West India Coastal Current (WICC), Lakshadweep Low (LL), Lakshadweep High (LH), South-West Monsoon Current (SMC), North-East Monsoon Current (NMC), SLD (Sri Lanka Dome) and East India Coastal Current (EICC). Findlater Jet is an atmospheric jet observed in the northern Arabian Sea during the summer monsoon. Processes associated with the summer monsoon are in red and those associated with the winter monsoon are in blue. WICC and EICC occur in both monsoons but differ in direction as indicated by the coloured arrows. Schematic adapted from Figure 1 in Luis & Kawamura (2004), Figure 1 in Peng et al. (2015) and Figure 1 in Vinayachandran et al. (2004).

The upper ocean circulation along the coasts of India is driven by seasonally reversing monsoon winds (Schott & McCreary, 2001). The large-scale monsoon currents flow eastward during summer (June to September, summer monsoon) and westward during winter (November to February, winter monsoon). Coastal currents, namely the East India Coastal Current (EICC) and the West India Coastal Current (WICC) (Shetye & Gouveia, 1998), link coastal circulation with the large-scale monsoon circulation. Saltier Arabian Sea water flows into the Bay of Bengal during the summer monsoon (Vinayachandran et al., 2018) and fresher Bay of Bengal water flows into the Arabian Sea during the winter monsoon.

During the summer (south-west) monsoon, the West India Coastal Current (WICC) flows towards the equator along the west coast of India (Shetye & Gouveia, 1998, Amol et al., 2014), which joins the South-West Monsoon Current (SMC). The latter turns around the southern tip of India and Sri Lanka and flows into the Bay of Bengal, connecting to the northward branch of the Sri Lanka Dome, which reverses direction at 10°N (Vinayachandran & Yamagata, 1998) (Figure 1). The East India Coastal Current (EICC) flows poleward during the summer monsoon (Mukherjee et al., 2014). During this period, coastal upwelling events of varying intensity occur along the west coast of India (Luis & Kawamura, 2004; Hood et al., 2017), with weak upwelling along the eastern coast of India, and the Andaman and Nicobar islands (Varkey et al. 1996; Vinayachandran et al., 2021).

During the winter (north-east) monsoon, the circulation along the east coast of India reverses direction to flow equatorward (EICC) (Shetye & Gouveia, 1998, Mukherjee et al., 2014), turns around Sri Lanka and the southern tip of India (Winter Monsoon Current) and flows poleward along the west coast (WICC) (Figure 1, Shetye & Gouveia, 1998; Shankar et al. 2002; Amol et al., 2014). Together with alongshore flow reversal during the winter monsoon, upwelling reverses into downwelling events within a 40 km wide band along the east coast of India (Varkey et al. 1996) and along the north-west coast of India (Luis & Kawamura, 2004).

The seasonal wind-driven circulation described above is modulated at several time-scales (Schott et al. 2009). The strong atmospheric-oceanographic coupling in the entire Indian Ocean makes its circulation sensitive to atmospheric intra-seasonal oscillations acting at time-scales ranging from a week (northward propagating precipitation anomalies) to 30-60 days (Madden-Julian Oscillation, Madden & Julian 1972). Inter-annual variability in the Indian Ocean circulation arises not only from El Niño Southern Oscillation, which leads to a year-long basin-scale warming after El Niño events, but also from Indian Ocean Dipole events, with cool (warm) and dry (wet) anomalies in the eastern (western) Indian Ocean in some years (Vinayachandran et al. 2009).

In such a complex oceanographic context with large to meso-scale structures, various levels of flow connectivity are to be expected along the vast Indian coastline, and it is difficult to a priori anticipate how biological filters such as spawning timing and PLD alter them. Identifying spatial scales at which demographic connectivity and/or gene flow between populations ceases to exist to give rise to biogeographic boundaries, is essential to inform marine resources and biodiversity management (Mertens et al., 2018).

In the current study, we aim to describe patterns of oceanographic connectivity focusing on the Indian coastline. To do so, we carried out Lagrangian transport simulations of neutrally buoyant larvae released along the study region’s coastline during the two major seasons – summer and winter monsoon for three years and across a wide range of PLD. Transport simulations were post-processed to build coastal connectivity networks for different combinations of season, year and PLD. The specific aims of our study were to (1) identify breaks in oceanographic connectivity and analyze their stability across years, seasons and PLD, (2) identify the most central locations within the connectivity network of the larger Indian coastline, and (3) examine the implications of these connectivity networks for biodiversity management.

## Methods

### Ocean circulation model

The HYbrid Coordinate Ocean Model (HYCOM, Chassignet et al., 2007) is an ocean general circulation model, which uses a combination of vertical coordinate systems (isopycnal, z-coordinate and sigma levels) to effectively simulate ocean circulation in three dimensions. The output from this circulation model is available for the global ocean as gridded values of horizontal velocities (eastward and northward), sea surface elevation, salinity and temperature at 1/12° spatial resolution, 3-hour temporal frequency and for 40 z-levels in depth. The model is forced by wind stress and fluxes of heat and freshwater at the surface. This model, however, does not include tidal flows, which would contribute to small-scale coastal processes. Particle transport in the vertical, which is particularly important in coastal areas with upwelling or downwelling, was accounted for after reconstructing vertical velocity values. To do so, the continuity equation of mass conservation was applied to horizontal velocities, and sea surface elevation data for the upper 27 depth levels from surface, i.e. down to 400 m deep. Beyond this depth, the coarse vertical resolution of HYCOM data can lead to anomalies in the derived vertical velocity values. Thus, the horizontal velocities obtained directly from HYCOM and the vertical velocity derived using the continuity equation was used to drive the particle tracking model from the free surface down to 400 m depth.

We used HYCOM output (GOFS3.0: HYCOM + NCODA Global 1/12° Reanalysis, GLBu0.08: expt_19.1 – https://www.hycom.org/dataserver/gofs-3pt0/reanalysis) for the spatial extent ∼30°N-10°S, 50°W-100°E and the 2008-2011 time period for particle tracking simulations spanning the winter (north-east) monsoon (November to February), and the summer (south-west) monsoon (June to September). These years were chosen based on availability of current data, and to capture the variation in ocean circulation presented by various states of the El Niño–Southern Oscillation (ENSO). While 2009-2010 represents an El Niño year, 2011 was a La Niña year (https://psl.noaa.gov/enso/past_events.html).

### Particle tracking model

Connectivity Modeling System (CMS) is an open-source, offline three-dimensionnal Lagrangian particle tracking model written in Fortran, which is compatible with HYCOM output. It works by interpolating velocity values to a given particle position using a fourth order Range-Kutta stepping scheme and advects the particle over a user-defined time-step (Paris et al., 2013).

In this study, we ran three dimensional particle tracking simulations, where larvae were simulated as neutrally buoyant particles, which passively drifted with ocean currents. Particles were released along the coastline of the Indian subcontinent (Indian coastline and parts of neighbouring Pakistan to the west and Bangladesh to the east), Sri Lanka, Lakshadweep islands and the Andaman and Nicobar islands. Particles were released every 5 km at 1 m below the surface, yielding a total of 2136 release locations along the length of the study area’s coastline. Particle release locations were created using the packages *‘raster’* (Hijmans, 2020), *‘rgeos’* (Bivand & Rundel, 2020), *‘sp’* (Pebesma & Bivand, 2005; Bivand et al., 2013) in R version 3.6.1 (R Core Team, 2019), and further edited in QGIS 3.14.0-Pi (QGIS.org, 2021).

The release frequency was one particle every three hours from each location between November to February (winter monsoon), and June to September (summer monsoon), summing up to about six million particles tracked during each monsoonal season. Particle positions were calculated using a time-step of 20 minutes, their coordinates were recorded every three hours, and each particle was tracked for a duration of 50 days after release. The tracking duration for particle trajectories was determined based on maximum values of pelagic larval duration reported for marine invertebrate taxa found in this region (Supplementary Information 1). For the purpose of analysis, each season was divided into 12 release bouts of ten days each to capture intra-seasonal variation in hydrodynamics that can influence particle trajectories.

### Connectivity networks

Coastal polygons, each of area ∼200 km^2^, including four release locations and numbering 528 in total, were created along the coast of the study area (Supplementary Information 2) using the packages *‘raster’* (Hijmans, 2020), *‘rgeos’* (Bivand & Rundel, 2020), *‘sp’* (Pebesma & Bivand, 2005; Bivand et al., 2013) in R version 3.6.1 (R Core Team, 2019), and further edited in QGIS 3.14.0-Pi (QGIS.org, 2021). Connectivity matrices of larval flux were built by calculating the proportion of particles released in an origin coastal polygon that successfully dispersed to a destination coastal polygon for each PLD ranging from 2 to 50 days, with an interval of 2 days. Considering all polygons as potential origin and destination sites for larvae, the size of a larval flux matrix was 528^2^ (i.e. 2,78,784 potential connections). Laval flux per unit surface was obtained by scaling larval flux values with the the area of the destination polygon (Figs 2a, 2b). Given the frequency distribution of PLD for marine invertebrates found in the Indian Ocean, connectivity matrices were averaged into four PLD classes for each spawning period – 2-4 days, 6-12 days, 14-20 days and 22-50 days (Supplementary Information 3).

**Figure 2.**
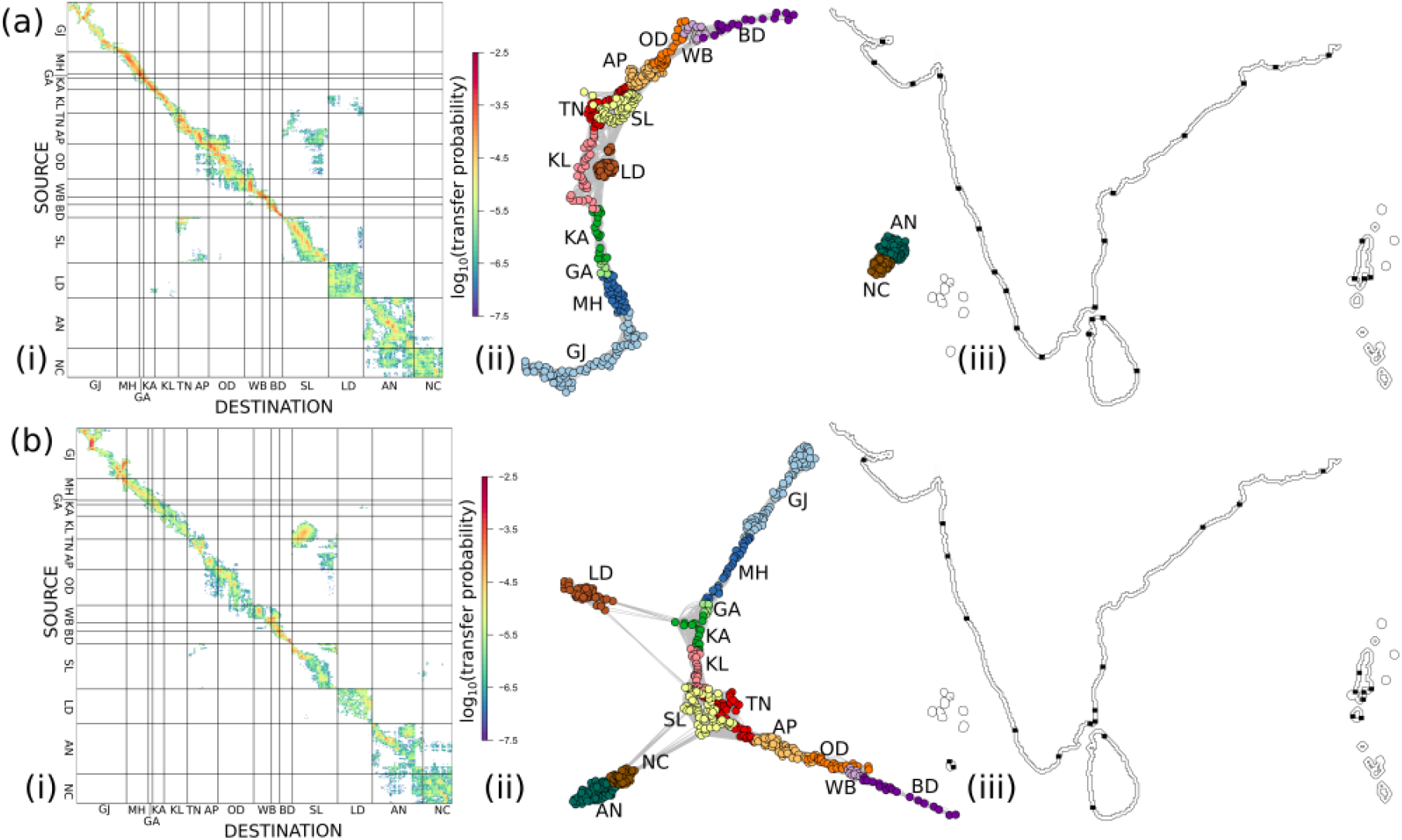
Schematic figure depicting the various stages in connectivity analysis. For illustrative purposes, the figures shown here are derived from a mean connectivity matrix across spawning periods and years for the PLD class 14-20 days, for (a) the winter monsoon and (b) the summer monsoon. For each monsoon, subfigures are (i) connectivity matrix represented as log_10_(transfer probability), (ii) connectivity network obtained from the raw transfer probability matrix and (iii) network community boundaries (black squares along the coastline) detected by applying the Infomap algorithm on the connectivity network. The codes presented in (i) and (ii) correspond to GJ – Gujarat, Diu and a section of Pakistan coast, MH – Daman and Maharashtra, GA – Goa, KA – Karnataka, KL – Kerala, TN – Tamil Nadu, AP – Andhra Pradesh, OD – Odisha, WB – West Bengal, BD – section of Bangladesh coast, SL – Sri Lanka, LD – Lakshadweep islands, AN – Andaman islands and NC – Nicobar islands.

In total, 288 connectivity matrices were built corresponding to the different PLD-classes, spawning periods, seasons and years (4 PLD-classes × 2 seasons × 12 spawning periods × 3 years). Each connectivity matrix defined a connectivity network, where nodes were coastal polygons and directed weighted edges between each pair of nodes were values of larval flux per unit surface between them (Figs 2a, 2b). Various metrics were used to describe the connectivity network – connectance (proportion of non-zero edges), number of singleton nodes (nodes not connected to any other node within the network) and number of weak subgraphs (group of nodes connected through paths in a single direction) or strong subgraphs (group of nodes connected through paths in both directions) based on the nature of connectivity.

### Community detection

Each of the 288 connectivity networks was individually processed to detect communities using the Infomap algorithm (Rosvall & Bergstrom, 2008) (Figs 2a, 2b) by applying the *‘cluster_infomap’* function in the *‘igraph’* package (Csardi & Nepusz, 2006) in R. This algorithm uses information-theoretic principles to define a community based on the ease of flow between network nodes. A community is detected when a random walker visits a set of nodes within the network more often than nodes outside it and is likely to get trapped between them (Rosvall & Bergstrom, 2008). The significance of a detected community was estimated using its coherence ratio – the proportion of particles that originate from nodes within a community and remain within the same community (Rossi et al., 2014). A coherence ratio greater than 0.5 was applied to detect well connected communities, and the location of community boundaries (referred to as community breaks henceforth) was detected. Community breaks were classified based on their frequency of occurrence across spawning periods within a PLD-class. Those occuring with a frequency greater than 50% were termed ‘frequent’ and those with a frequency greater than 90% were termed ‘highly consistent’.

### Centrality measures

For centrality measures using shortest path calculations, larval flux values were transformed into distance between nodes by applying a log(1/x) transformation (Costa et al., 2017). The role and importance of each node (coastal polygon) in the connectivity network (PLD-class × season × spawning period) was assessed using six different metrics. Node degree (number of non-zero edges from a node) and strength (sum of weight of edges from a node) were used to rank nodes according to their overall influence on the network (Dubois et al., 2016). Betweenness centrality, which measures the number of shortest paths between node pairs that pass through a given node (Freeman, 1977), was used to identify nodes which control transport within the connectivity network. Clustering coefficient, the proportion of realized directed triangles (three nodes connected in all possible ways) between a focal node and its immediately connected nodes (Fagiolo, 2007), was used to measure the propensity of a node to cluster with its neighbours leading to redundancy within a network. Bridging centrality, a product of betweenness centrality and bridging coefficient (node ranking based on location between dense network sub-graphs) (Hwang, 2008), was used to identify nodes which control connections between network communities. Closeness centrality, the inverse of sum of pairwise shortest distances between a given node and all other nodes in a network, was used to identify nodes that can independently communicate with different regions within strong subgraphs in a network (Freeman, 1978).

Centrality measures were calculated using the tattoo toolbox (https://github.com/costaandrea/TATTOO, Costa, 2017) in MATLAB R2019b.

### Statistical tests

The significance of variation in the distribution of connectance and the number of singleton nodes, weak and strong subgraphs, and Infomap communities with >1 membership, with PLD-class, year and season was evaluated using non-parametric Kolmogorov-Smirnov test in R version 3.6.1 (R Core Team, 2019).

## Results

There was no significant variation in most connectivity network characteristics between years, while their variation between spawning periods within a year was found to be large (Supplementary Information 4). Based on these results, variability of all metrics was estimated by pooling 36 connectivity matrices across years (3 years × 12 spawning periods) for each PLD-class and season.

### Network descriptors

Connectance was low along the Indian coastline, irrespective of the PLD-class and season, with values ranging from 0.5% to 3.4%. During both the seasons, connectance showed the greatest significant difference between PLD less than 20 days and PLD greater than 20 days (Figure 3). Despite the low connectance, there were less than ∼30% singleton nodes across PLD-classes and seasons, indicating that transport further than 5 km along the coast (minimum distance between release locations in adjacent coastal polygons) was common (Figure 3). The number of singleton nodes increased till a PLD of 20 days, indicating that local connections with neighbouring coastal polygons at shorter PLDs disappeared at intermediate PLDs as particles dispersed away from the coast. Except for the PLD-class 6-12 days, the number of singleton nodes was not significantly different between seasons (Supplementary Material 3).

**Figure 3.**
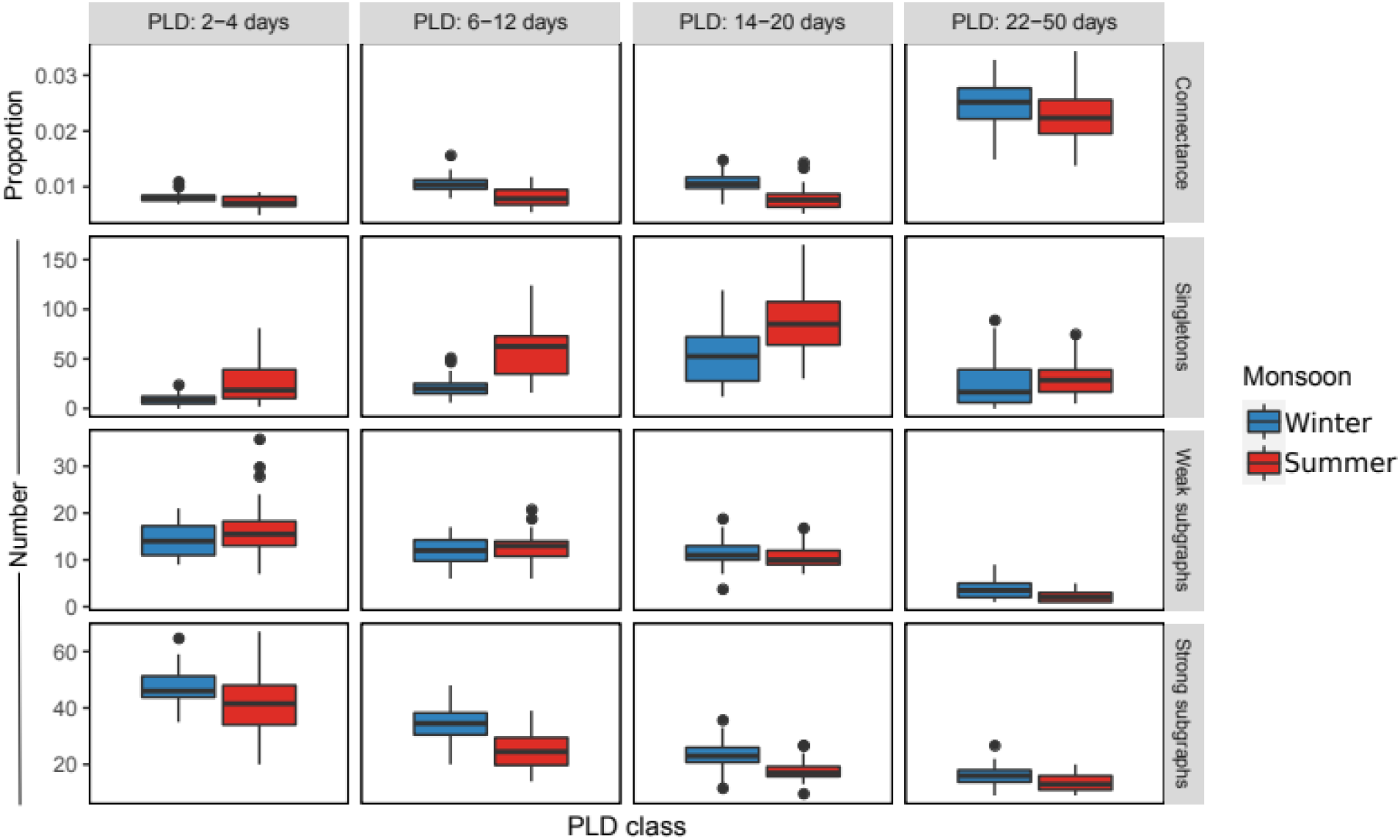
Summary characteristics of connectivity networks.

The number of weak subgraphs largely remained unchanged for PLD less than 20 days (ranging from 4 to 36), while the number of strong subgraphs significantly decreased between the PLD less than 6 days (ranging from 20 to 67) and the PLD greater than 20 days (ranging from 9 to 27) (Supplementary Material 4). This indicates that there was an increase in the number of bi-directional connections within subgraphs with increasing PLD. This densification of subgraphs, for PLD up to 20 days, was more pronounced in the summer monsoon as compared to the winter monsoon (Supplementary Information 4).

Put together, a significant increase in the connectance for PLD greater than 20 days indicates that a longer transport duration enabled greater transport distance along the coast and islands, leading to a decrease in the number of isolated singleton nodes and an increase in subgraph size. A concomitant decrease in the number of weak and strong subgraphs for PLD greater than 20 days indicates the formation of more densely connected subgraphs.

### Infomap communities

Similar to the trend observed with strong subgraphs, the number of Infomap communities (communities hereafter) decreased with an increase in PLD (Figure 4a). Except for the PLD-class 6-12 days, the number of communities detected for the same PLD-class was not significantly different between seasons (Supplementary Material 4). The coherence ratio of the detected communities was significantly different between seasons (Supplementary Material 4) and decreased with increasing PLD as seen from the increase in variance within a PLD-class (Figure 4b). The decrease in coherence ratio indicates a decrease in the flux of particles circulating within a community through losses outside it.

**Figure 4.**
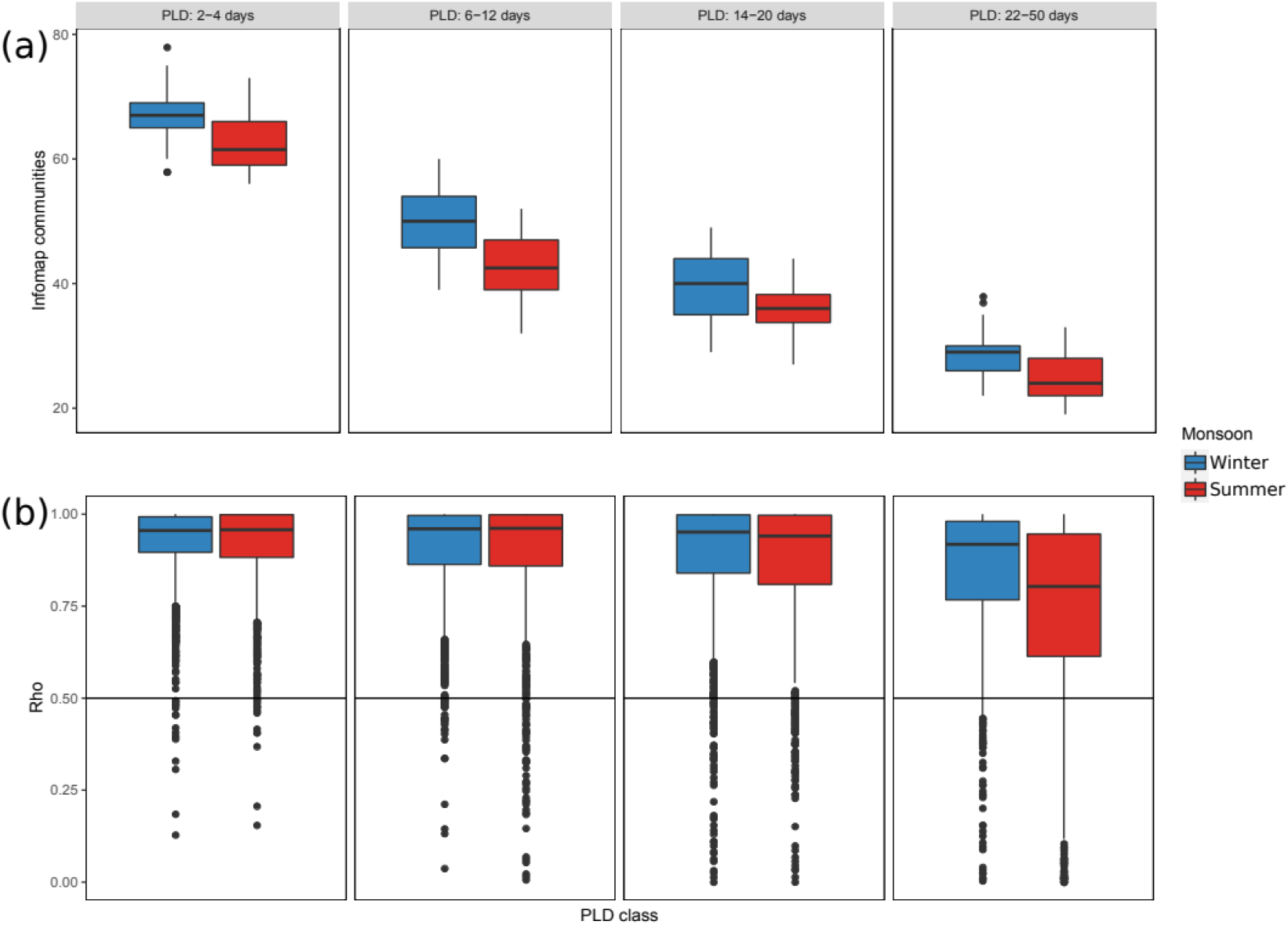
(a) Number of Infomap communities and (b) their coherence ratio.

The location of community breaks varied with PLD, season and spawning period (indicated by the variation in the consistency of breaks within a PLD-class). However, a few locations frequently appeared as community breaks across spawning periods (>50% frequency), with some occuring consistently with a frequency greater than 90% (Figure 5). These community breaks were found along the north-west coast of mainland India (south of the Gulf of Khambhat 21°N, and Gulf of Kutch), Palk Strait (a narrow channel separating the Indian landmass from Sri Lanka) and north-east India, across both seasons (Figure 5). In the summer monsoon, a consistent community break appeared at the southern tip of India for all PLDs. Additional breaks also appeared along the west and east coast of India for PLD less than 20 days, but were not consistent across season and PLD.

**Figure 5.**
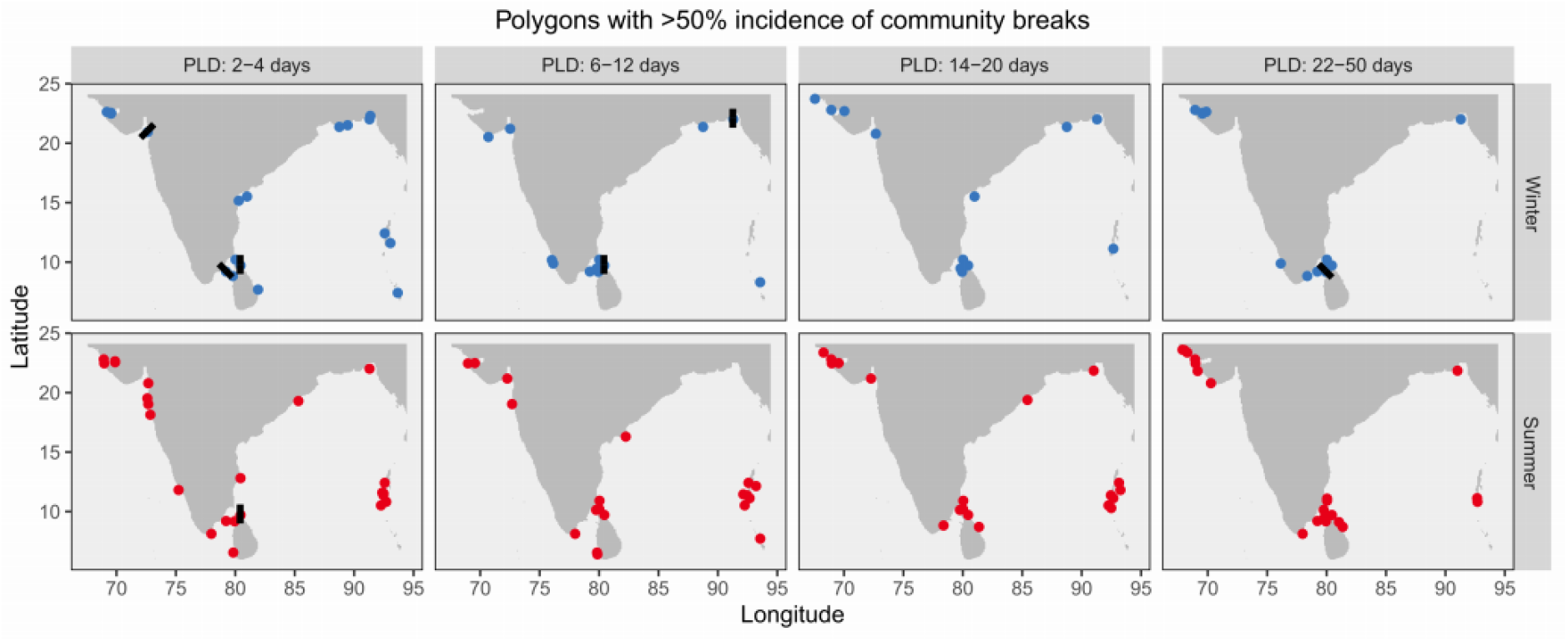
Distribution of Infomap community breaks. Blue and red circles indicate breaks with frequency >50% across spawning bouts for the winter and summer monsoon respectively. Black bars indicate breaks with frequency >90% across spawning bouts.

### Node descriptors

As observed with connectance, node degree increased with PLD as a longer transport duration promoted variability in transport from the same release location (Figure 6). However, the increase in node degree was not uniform in the study region, and was observed to be higher in the east coast of India and the Lakshadweep islands during the winter monsoon, southern tip of India during the summer monsoon, and Sri Lanka and the Andaman and Nicobar islands during both the seasons (Figure 6). Another consequence of transport over longer durations was the loss of particles from the coastal zone as observed from a decrease in the node strength across a majority of nodes across seasons (Figure 6). Node strength decreased more gradually with increasing PLD, and nodes with high values were more evenly distributed in space in the winter monsoon as compared to the summer monsoon (Figure 6).

**Figure 6.**
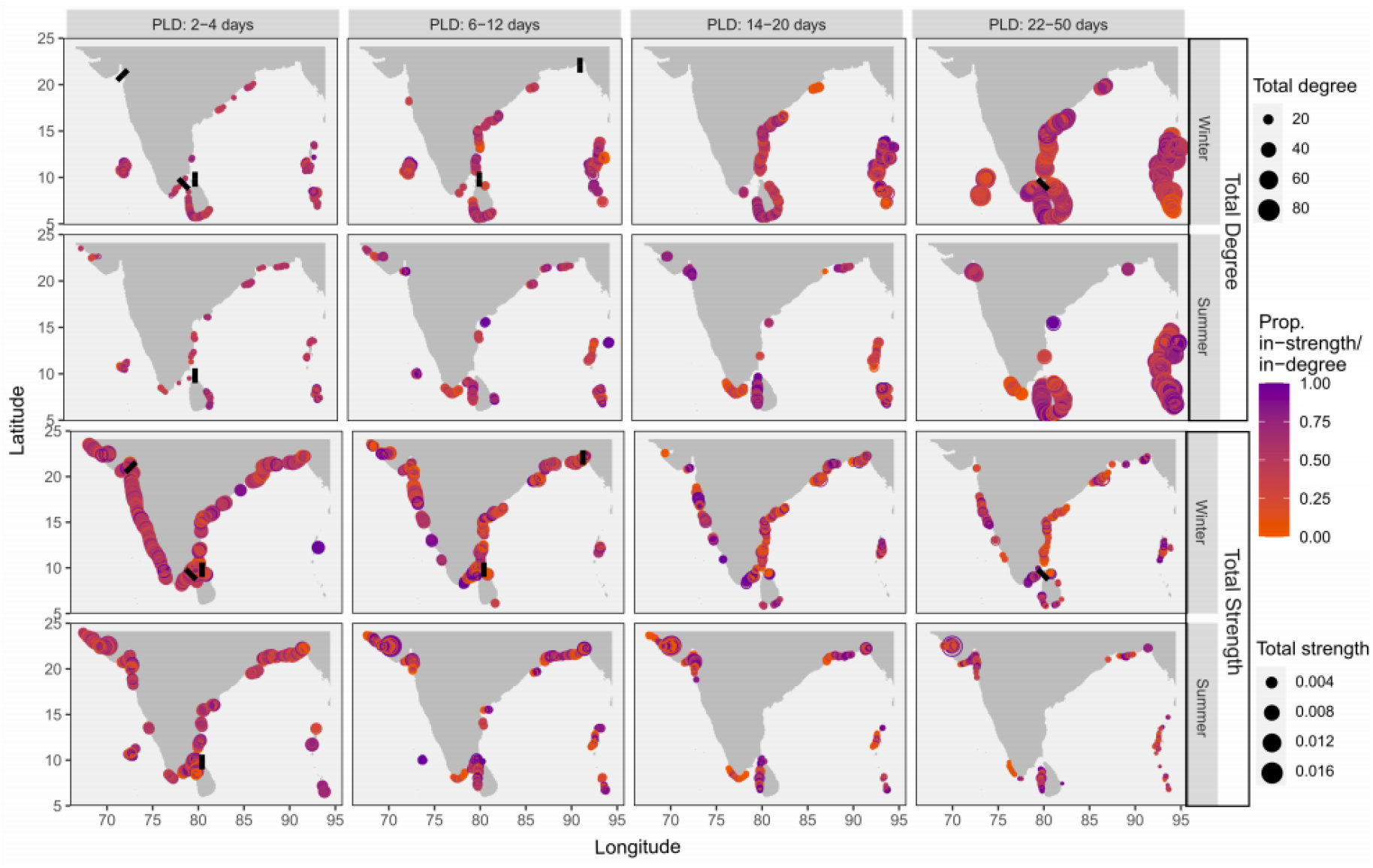
Spatial distribution of total strength (sum of incoming and outgoing edge weights) and degree (sum of number of incoming and outgoing edges) for nodes which have strength/degree values greater than the median atleast 75% times across spawning bouts within a given PLD-class. The colour of a node indicates the magnitude of in-strength/degree, ranging from orange (source) to blue (sink). At each node, filled circles represent the median and the concentric open circles represent the 75^th^ quantile of degree/strength. Infomap community breaks with >90% incidence are shown as black bars.

The number of nodes with high values of centrality measures reduced drastically for PLD greater than 6 days with spatial disparity based on the season. For the shortest PLD, betweeness centrality and bridging centrality were higher and more uniformly distributed along the west coast of India as compared the east coast, while the opposite pattern was observed for closeness centrality. For this PLD range, limited transport led to high clustering coefficient along large extents of the Indian coastline in both seasons (Figure 7).

**Figure 7.**
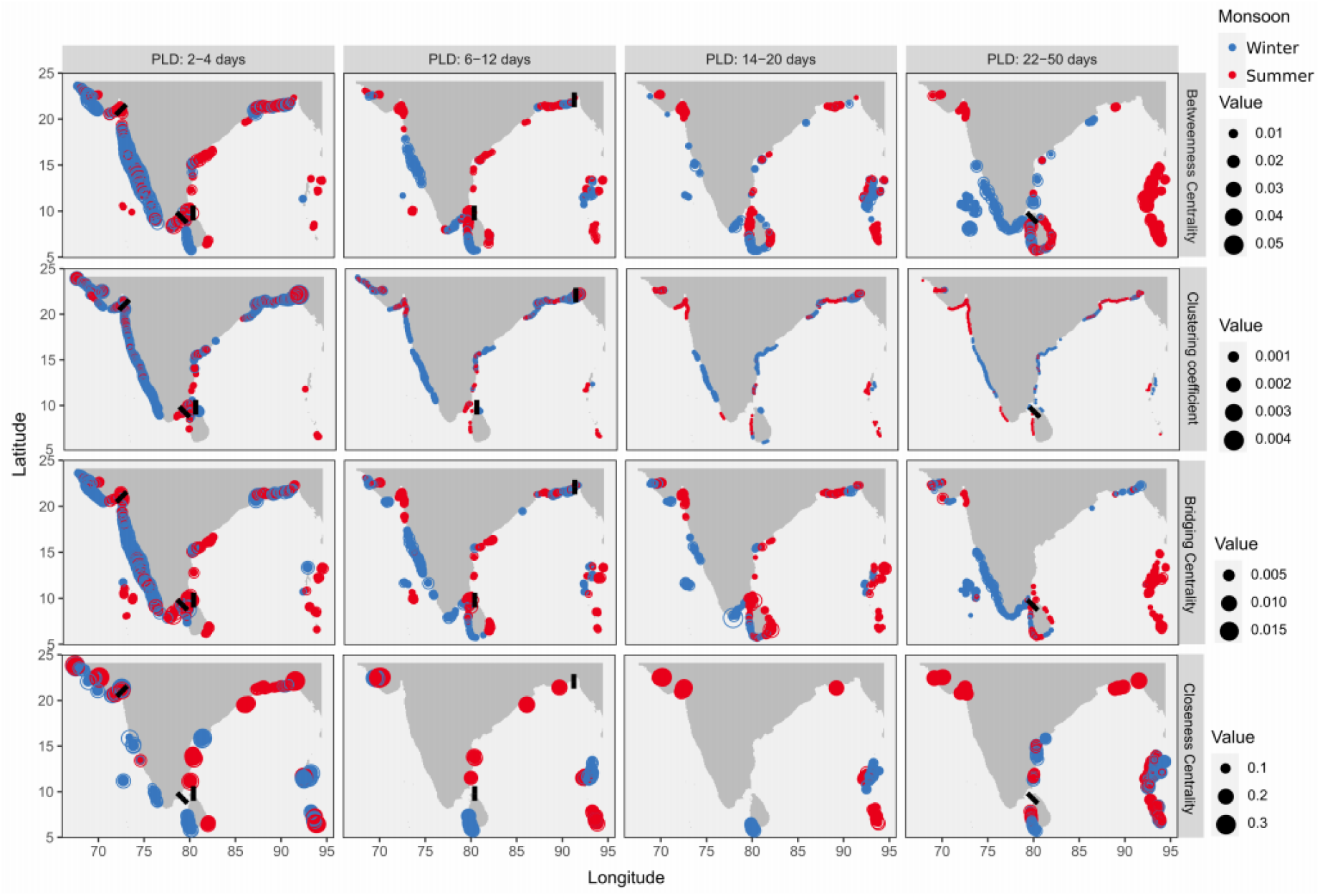
Spatial distribution of important nodes based on measures of network centrality. The nodes displayed on the map have centrality values greater than the median atleast 75% times across spawning bouts within a given PLD-class. At each node, filled circles represent the median and the concentric open circles represent the 75^th^ quantile of the centrality measure. Infomap community breaks with >90% incidence are shown as black bars.

High betweenness and bridging centrality values, depicting nodes controlling larval transfer within a network, were consistently observed in regions along the west coast, south-east coast and western Sri Lanka during the winter monsoon, while in the summer monsoon they occurred in Sri Lanka, north-west coast and the Andaman and Nicobar islands. Both the island groups showed high betweenness and bridging centrality, but did not show consistent patterns across PLD and season. Nodes with high clustering coefficient, indicating node redundancy within a network, were observed in central east and west coasts during the winter monsoon, and the north-west coast and the Bangladesh coastline during the summer monsoon. Nodes with high closeness centrality varied with season and PLD, but the Gujarat and Bangladesh coasts displayed consistently high closeness values, thus facilitating direct larval transfer within the network communities (Figure 7).

## Discussion

This is one of the first broad-scale studies to use ocean flow data and particle tracking models to describe patterns of coastal connectivity along the coastline of the Indian subcontinent and the adjacent island groups. We found that properties of connectivity networks exhibited intra-season and inter-season variability showing differences with pelagic larval duration (PLD), but were comparable across years. Though values of connectance were generally low within the coastal network, there was an increase in the size and bidirectionality of network subgraphs with increasing PLD, resulting in fewer and larger well connected coastal communities.

Despite seasonal differences in the location of community breaks, particularly for PLD less 20 days, four disconnected provinces along the mainland coast were identified across PLD and seasons put together. From west to east along the coastline of the Indian subcontinent, the first province extended across the Gulf of Khambat and the coast of Gujarat (southern boundary at 21°N), the second province was found along the west coast of India (∼21°N to 8°N), the third province observed in the summer monsoon extended from the southern tip of India to the Palk strait (Gulf of Mannar, between Sri Lanka and mainland India) and the fourth province was observed along the east coast of India (∼11°N up to 22°N). Within each of these provinces, a few sites displayed notably higher centrality values than others, but their importance varied based on season and PLD.

### Oceanographic drivers of connectivity

A striking feature of the upper ocean circulation along the Indian coastline is the seasonal reversal of coastal currents, which are anti-clockwise in the summer monsoon and clockwise in the winter monsoon. Despite the latter, three of the four coastal provinces delineated in our study remained stable across seasons. Such stability is consistent with two persistent hydrodynamic barriers – one north of 20-21°N in the Arabian Sea (Luis & Kawamura, 2004) and the other associated with a deviation in coastal circulation around Sri Lanka near the southern tip of the Indian coastline (Schott & McCreary, 2001).

However, some hydrodynamic barriers and connectivity network descriptors exhibited intra-seasonal variability due to an interaction of spawning timing within a reproductive season with intra-seasonal flow variability. This flow variability is largely driven by atmospheric intra-seasonal oscillations in the Indian Ocean, resulting in short-lived upwelling/downwelling events along the Indian coastline (Varkey et al., 1996; Luis & Kawamura, 2004; Durand et al., 2009). These oscillations span over a wide range of time-scales ranging from a week to 60 days (Madden & Julian, 1972; Fu et al., 2003), which likely explains why intra-seasonal variability was not averaged out across the PLD range of 2 to 50 days considered in the present study.

Inter-annual atmospheric oscillations, such as the El Niño–Southern Oscillation and the Indian Ocean Dipole, are also known to alter Indian Ocean climatology and large scale circulation (reviewed in Schott et al. 2009). Interestingly, though the resolution of a Global Ocean Circulation Model such as HYCOM is more suitable to describe inter-annual variation as compared to short lived atmospheric events, we find that intra-seasonal variability was greater than inter-annual variability for all descriptors of network connectivity. These results advocate for fine-tuning connectivity studies in the future to include a better description of short-lived atmospheric processes and to down-scale flow modelling within each of the provinces delineated in the present study.

### Biotic filters of connectivity and biogeographic boundaries

Biological traits related to spawning season and PLD determine the time-dependent ocean flow scenarios that larvae are exposed to and act as filters over the connectivity patterns that are eventually realized in a given region. By combining empirical information on spawning season and PLD range of the species of interest (Supplementary Information 1) with connectivity descriptors from the present study, it is possible to project taxa-specific predictions of connectivity breaks.

For instance, in the case of both anthozoans (PLD: <7 days, likely summer spawning) and holothurians (PLD = 12-28 days, likely summer spawning), important connectivity breaks would be predicted to occur around the southern tip of India, Palk Strait, south-west Sri Lanka and ∼20-21°N on the west coast of India. For several crustaceans (PLD = 8-43 days, likely winter spawning) and gastropods (PLD = 12-46 days, likely winter spawning), connectivity breaks would be predicted to occur around the Gulf of Khambat, Palk Strait and north-east Indian coastline. Finally, for taxonomic groups such as bivalves and non-holothurian echinoderms, connectivity patterns are not generalizable across species because of a large variability in their spawning period (bivalves) or PLD (non-holothurian echinoderms).

The occurrence of genetic breaks in connectivity has only been tested in a few studies of marine invertebrates along the Indian coastline, such as in the Indian prawn (the crustacean *Penaeus indicus*, PLD: 12-25 days, spawning: October-April) (Sajeela et al., 2019), intertidal periwinkles (the gastropod *Littoraria* species, PLD: 21-70 days, spawning: throughout year; *Echinolittorina* species, PLD: 21-28 days, spawning: March-June) (Bharti, 2019) and the Asian green mussel (the bivalve *Perna viridis,* PLD = 17.5 days, spawning period: contrasting between coasts) (Divya et al., 2020). Across these PLD and spawning period combinations, the connectivity break consistently predicted around southern India coincided with patterns of population genetic connectivity (Figure 8a). For *Penaeus indicus* and *Echinolittorina malaccana,* genetic breaks also coincided with the predicted connectivity break around 21°N on the west coast of India. However, in many of these studies, the populations are separated by large gaps in sampling, which makes it difficult to identify all the genetic breaks and, more importantly, to differentiate between the role of isolation by distance versus dispersal barriers in shaping the observed patterns (Audzijonyte & Vrijenhoek, 2010).

**Figure 8.**
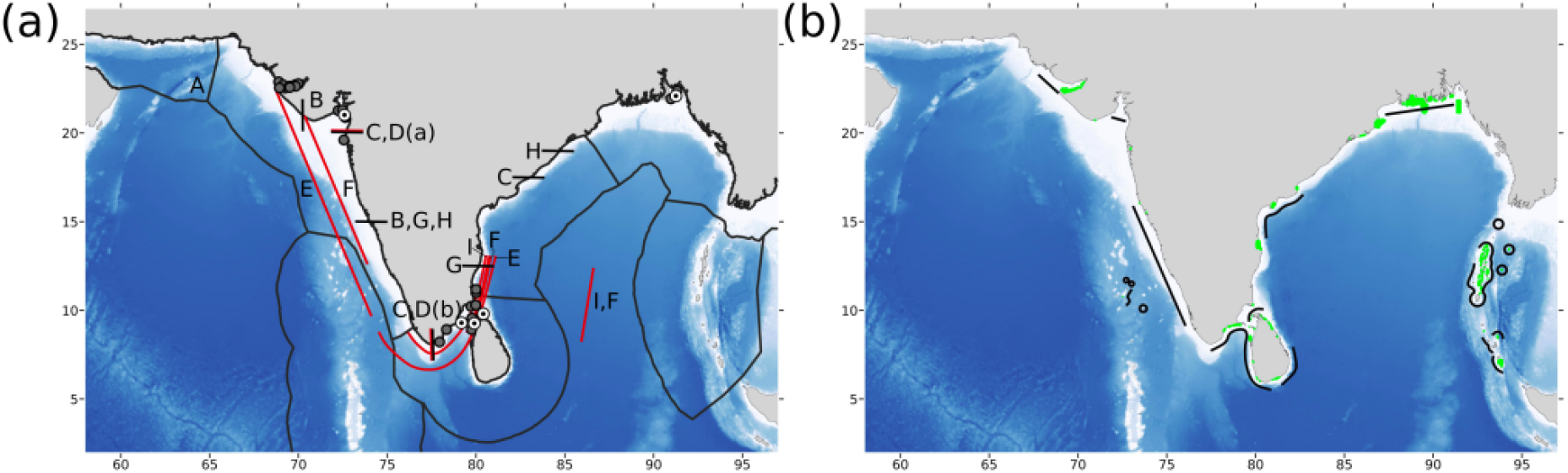
(a) Biogeographic boundaries (black lines) and genetic breaks (red lines) observed for marine species along the Indian coastline – A. Marine ecoregions along the Indian coastline (Spalding et al., 2007), B. Boundaries of Ecological Marine Units (Sayre et al., 2017), C. Subdivision of marine ecoregions based on distribution of gastropods, bivalves and polychaetes (Sivadas & Ingole, 2016), D. Genetic breaks observed for the intertidal snails (a) *Echinolittorina malaccana* and (b) *Littoraria strigata* (Bharti, 2019), E. Extent across which genetic breaks are observed for the marine fish *Rachycentron canadum* (Divya et al., 2017), Extent across which genetic breaks are observed for the shrimp species *Penaeus indicus* (Sajeela et al., 2019); G. Biogeographic divisions based on distribution of bivalve species (Sarkar et al., 2017); H. Boundaries of distribution zones defined for pelagic fisheries (Pillai et al., 2007) and I. Extent across which genetic breaks are observed for the mussel *Perna viridis* (Divya et al., 2020). Community breaks identified from this study are depicted using grey (>50% frequency across spawning periods) and concentric black (>90% frequency across spawning periods) circles. (b) Areas highlighted in green indicate marine protected areas (outlines exaggerated for visibility) and areas highlighted using black lines indicate regions observed to have high betweenness and bridging centrality from this study.

The consistent community breaks predicted by our larval transport models in the Palk Strait in both seasons and in the southern tip of India during the summer monsoon, correspond to the separation of southern India and Sri Lanka from the east and west coasts suggested by the Spalding et al’s (2007) classification of ecoregions along the world’s coastlines (Figure 8A). This scheme uses published studies from multiple taxonomic groups and oceanographic processes to define biogeographic boundaries, where ecoregions represent areas showing similarities in species composition, which may be driven by unique oceanographic and geomorphological features. The distinct biogeography of the east and west coasts of India is also suggested by Ecological Marine Units defined by environmental parameters (Sayre et al., 2017), and patterns of community composition in marine invertebrates along the Indian coastline (Sivadas & Ingole, 2016; Sarkar et al., 2017). (Figure 8a)

The connectivity break south of the Gulf of Khambat, separates the linear coast of western India from the Gujarat, and is seen across at least half of all spawning periods for both seasons and PLD less than 20 days. This corresponds to the classification of an ecological marine unit extending across the Gulf of Oman to the Oman coastline (Sayre et al., 2017). This region is characterized by seasonal variation in salinity, with the formation of the Arabian Sea High Salinity Water Mass (Kumar & Prasad, 1999) in the winter monsoon and upwelling during the south-west monsoon, which influences productivity and zooplankton biomass (Madhupratap et al., 1996; Madhupratap et al., 2001). This indicates that ocean flow along with environmental features might drive patterns of biogeography in the northern Arabian Sea.

However, it is important to highlight the limitations of large-scale transport simulations of neutrally buoyant particles using PLD and spawning season alone, which may lead to an incomplete understanding of taxa-specific connectivity breaks. Firstly, larval transport is influenced by species-specific variation in buoyancy and swimming behaviour, which can deviate from predictions for neutrally buoyant larvae (Guizien et al., 2006; Robins et al., 2013), resulting in different connectivity patterns (Le Corre et al. 2018; Blanco et al. 2019). Secondly, large-scale ocean flow simulations, with a spatial resolution of ∼10 km, are inadequate to describe meso-scale variability of coastal circulation. Such fine-scale variability has been shown to shape the connectivity of coastal populations in fragmented and spatially complex habitats (Padrón et al. 2018; Frys et al., 2020). Thirdly, filters acting pre- and post-transport (sensu Pineda et al. 2007), which are not accounted for in the current study, are likely to explain additional breaks at the level of demographic or genetic connectivity. Finally, simulations of neutrally buoyant larval transport are not likely to identify connectivity breaks driven by ocean fronts exhibiting sharp differences in temperature and salinity (Sayre et al., 2017) (Figure 8a). This explains why some of the observed biogeographic boundaries, such as those reported in the central east and west coasts for bivalves (Sarkar et al., 2017) and pelagic fishes (Pillai et al., 2007) (Figure 8a), were not predicted as community breaks in our simulations.

### Implications for biodiversity conservation and management

The community breaks identified from our study mark the scale and extent of metapopulation connectivity along the Indian coastline, and can be used to guide the management of marine biodiversity. The broad connectivity provinces from our study include the coast of Gujarat, the linear extent of the west coast, southern India and the east coast of India. Currently, 24 National Parks and Wildlife Sanctuaries have been identified as Marine Protected Areas (MPA) on the mainland coast of India (Figure 8b), which are located within each of the provinces, with the exception of the linear west coast. Some of these MPAs are located in areas with recurrent connectivity breaks such as the Gulf of Mannar, Palk Strait, Andaman and Nicobar islands and the Gulf of Kutch. In these regions with complex coastal bathymetry, where connectivity breaks were detected at small spatial distances, community detection can be better defined by increasing the spatial resolution of ocean flow and the distribution of release locations (Briton et al. 2018). Spatially and temporally refined transport studies are key in accounting for the effect of small-scale ocean features on flow connectivity at the resolution of coastal populations and local habitat distribution. This defines the relevant spatial scale for assessing vulnerability of coastal populations and designing protection measures (Guizien et al. 2012, 2014).

The use of centrality measures has been suggested to guide coordinated management at the network level in several studies (Andrello et al. 2013). We find that some areas acting as stepping stones of connectivity (indicated by high betweenness centrality), such as the Gulf of Kutch in the north-west, Gulf of Mannar in the south-east and Sunderbans in the north-east of India, are already under the MPA scheme (Figure 8b). Designation of additional areas for protection, especially along the west coast of India, can improve the existing MPA network. However, within each of the predicted connectivity provinces, the importance of centrality measures differs with season and PLD, indicating that temporally adaptive management measures specific to local biodiversity might be necessary.

To conclude, the present study highlights the importance of intra-season variability in driving patterns of connectivity and provides a coarse prediction of connectivity provinces that fulfill a gap in marine biogeography studies from this region. Our results can guide the spatial scales at which future biophysical models should be set up. Based on the frequency of community breaks, we advocate for refining transport studies based on flow modelling at the appropriate resolution along the Gujarat coastline, around the southern tip of India and Sri Lanka, over two large areas extending along the western and eastern coasts of the Indian subcontinent, and within each of the island groups. Our findings of seasonal and taxa-specific variation in areas exhibiting a large influence on connectivity patterns can inform regional biodiversity management along the Indian coastline.

## Acknowledgements

This work was supported by the Department of Biotechnology, Government of India (BT/PR15704/AAQ/3/758/2015). Collaborative work was facilitated by an EMBO Short-Term Fellowship (STF 7321) awarded to DKB to visit Observatoire Océnaologique de Banyuls Sur Mer (UPMC/CNRS), France. Research fellowship to DKB was awarded by the Council of Scientific and Industrial Research, Government of India (09/079(2450)/2011-EMR-I). DKB is grateful to Aarti Krishnamoorthy, Aditya Dharapuram and Lakshmi Prasad Natarajan for offering technical support for running the particle tracking simulations.

## Data availability statement

The HYCOM output of gridded horizontal velocities is open-source and available for download from https://www.hycom.org/dataserver/gofs-3pt0/reanalysis. R and MATLAB scripts used for analyzing simulated particle trajectories will be made available at https://github.com/bhartidk/larval_dispersal.

