## Supporting Information Appendix S1 for "Connectivity networks and delineation of distinct coastal provinces along the Indian coastline using large-scale Lagrangian transport simulations"

**Appendix 1.** Estimates of pelagic larval duration (PLD) for marine invertebrates found along the Indian coastline.

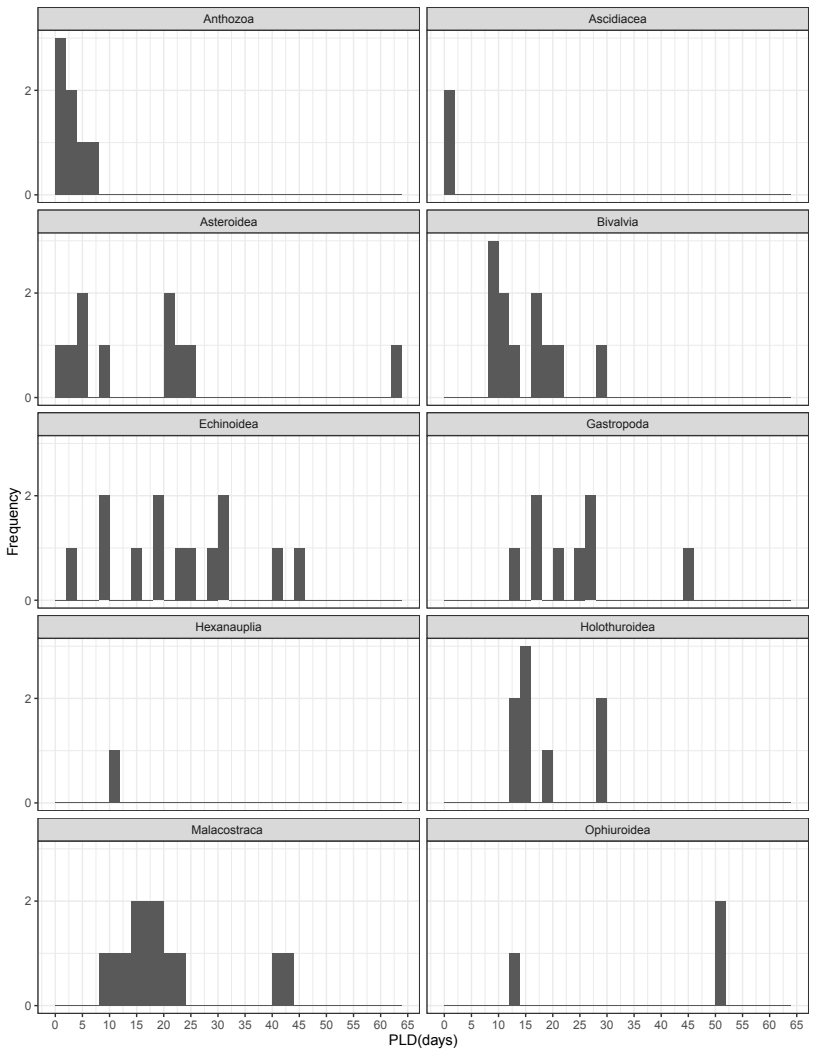

**Figure S1.1.** Distribution of PLD across marine invertebrates found in the coastal Indian Ocean by taxonomic class. This histogram omits one record of PLD 240 days from *Panulirus homarus* (Class Malacostraca).

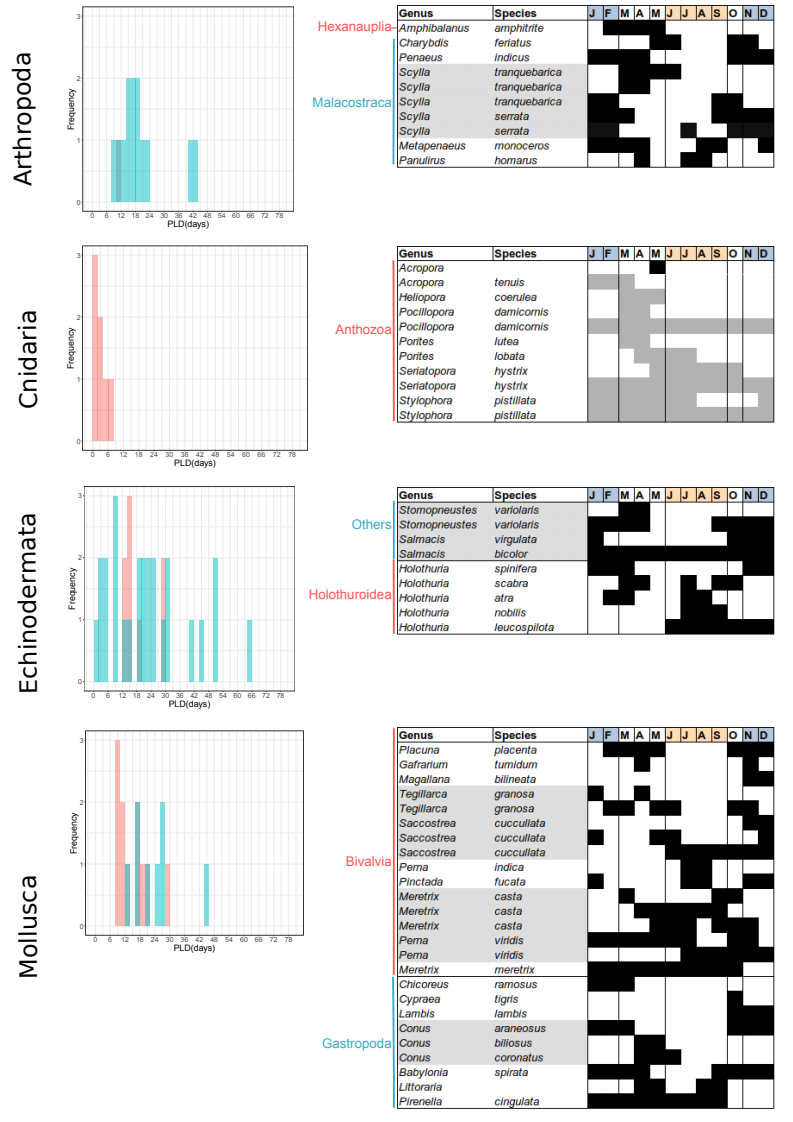

**Figure S1.2.** Distribution of PLD for marine invertebrates species found in the Indian Ocean region along with spawning period. Cells in black indicate spawning period reported from the Indian coastline, while grey is from neighbouring regions (refer to Table S1.1).

**Table S1.1.** List of marine invertebrate species with potential distribution in the Indian Ocean coast for which PLD and spawning period information was compiled.

| **Genus** | **Species** | **Phylum** | **Class** | **Mean PLD** | **PLD Reference** | **Spawn time** | **Spawn location** | **Spawn Reference** |
| --- | --- | --- | --- | --- | --- | --- | --- | --- |
| *Amphibalanus* | *amphitrite* | Arthropoda | Hexanauplia | 10 | Anil & Kurian, 1996; Anil, Desai & Khandeparkar, 2001 | Higher larval abundance during pre-monsoon February-May | Mandovi and Zuari estuaries, Goa | Gaonkar et al., 2012 |
| *Charybdis* | *feriatus* | Arthropoda | Malacostraca | 22.5 | Josileen, 2011 | Continuous breeders, peak in reproductively mature and berried females between October-November and May-June | Landing in Mangalore fisheries harbour (catch from north Kerala to southern Maharashtra) | Dineshbabu, 2011 |
| *Diogenes* | *miles* | Arthropoda | Malacostraca | 8.5 | Sankolli & Shenoy, 1993 |  |  |  |
| *Elamenopsis* | *demeloi* | Arthropoda | Malacostraca | 10 | Kakati, 1988 |  |  |  |
| *Exhippolysmata* | *ensirostris* | Arthropoda | Malacostraca | 43 | Bensam & Kartha, 1965; Kagwade, 1984; Pillai, 1974 |  |  |  |
| *Metapenaeus* | *moyebi* | Arthropoda | Malacostraca | 17.25 | Nandakumar et al., 1989 |  |  |  |
| *Metapenaeus* | *monoceros* | Arthropoda | Malacostraca | 19.15 | Mohamed, 1979; Nandakumar, 2001 | Year around, peaks from December-April and August-September | Individuals obtained from trawl landing, Kochi | Nandakumar, 2001 |
| *Metopograpsus* | *latifrons* | Arthropoda | Malacostraca | 18.5 | Kakati, 1972 |  |  |  |
| *Ocypode* | *ceratophthalmus* | Arthropoda | Malacostraca | 40 | Kakati, 2005 | Congener *Ocypode rotundata* breeds from March-October with a peak in June | Qeshm Island, Persian Gulf | Naderi et al., 2018 |
| *Ozius* | *rugulosus* | Arthropoda | Malacostraca | 20 | Kakati & Nayak, 1977 |  |  |  |
| *Panulirus* | *homarus* | Arthropoda | Malacostraca | 240 | Berry, 1974; Booth & Phillips, 1994 | Likely throughout the year, possible peaks in April, July-August | Individuals obtained from Chennai | Vijayakumaran et al., 2005 |
| *Penaeus* | *monodon* | Arthropoda | Malacostraca | 14 | Duda & Palumbi, 1999 | Congener *Penaeus indicus* spawns from October-April | Individuals obtained from Kochi fisheries catch | Rao, 1964 |
| *Penaeus* | *semisulcatus* | Arthropoda | Malacostraca | 12.6 | Devarajan et al., 1979 | Can continuously spawn for as long as 3 months under laboratory conditions with different treatments | Most likely hatchery raised individuals from Palk Bay | Radhakrishnan et al., 2000 |
| *Scylla* | *tranquebarica* | Arthropoda | Malacostraca | 17 | Maheswarudu et al., 2007 | Peak breeding from March-June^1^; September-February^2^; March-April and September-October^3^ | Chilika lagoon^1^, Kochi^2^, Pulicat lake^3^ | Mohanty et al., 2006 |
| *Scylla* | *serrata* | Arthropoda | Malacostraca | 15.35 | Hamasaki, 2003 | Continuous breeding, with peaks in December-March, September-November^1^; October-February; peaks in July and January^2^ | Karwar^1^; Kochi^2^ | Prasad & Neelakantan, 1989^1^; Thariyan, 1988^2^ |
| *Didemnum* | *molle* | Chordata | Ascidiacea | 0.08 | Olson, 1983; Olson, 1985 |  |  |  |
| *Phallusia* | *nigra* | Chordata | Ascidiacea | 0.8 | Lambert, 2002 |  |  |  |
| *Acropora* | *tenuis* | Cnidaria | Anthozoa | 7 | Nishikawa, Katoh & Kazuhiko, 2003 | Mature gametes of other Acropora species in March^1^, October-April with peak in January-March^2^ | Islands around Tuticorin^1^, Mombasa Lagoon Kenya^2^ | Raj & Edwards, 2010^1^, Mangubhai, 2008^2^ |
| *Heliopora* | *coerulea* | Cnidaria | Anthozoa | 0.25 | Harii et al., 2002; Harii & Kayanne, 2003 | March-May | Bolinao, Philippines | Vicentuan et al., 2008 |
| *Montipora* | *capitata* | Cnidaria | Anthozoa | 3 | Conception et al. unpubl. |  |  |  |
| *Pocillopora* | *damicornis* | Cnidaria | Anthozoa | 1 | Harii et al., 2002 | Continuous breeding season^1^. Spawning inferred in March-April, a month after highest summer temperature^2^; Continuous spawning throughout year, sometimes continuously for 3 months^3^ | Durban, South Africa^1^,^2^; Chonburi, Thailand^3^ | Masse et al., 2012^1.2^; Kuanui et al. , 2008^3^ |
| *Porites* | *lobata* | Cnidaria | Anthozoa | 3 | Selkoe et al., 2014 | Congener Porites lutea spawns from March-April^1^; *Porites lobata* is generally observed to spawn in summer or spring months across range | Singapore^1^ | Stoddart et al., 2012 and references therein |
| *Seriatopora* | *hystrix* | Cnidaria | Anthozoa | 1 | Maier et al., 2005 | Peak spawning from May-October in Okinawa, spawning duration may vary with latitude; In Phillipines it is year round | Okinawa, Japan | Prasetia et al., 2017 and references therein |
| *Stylophora* | *pistillata* | Cnidaria | Anthozoa | 4.5 | Nishikawa, Katoh & Kazuhiko, 2003 | December-July in Eliat; Throughout year in Palau; Possible relationship with latitude and temperature indicated | Gulf of Eliat, Red Sea | Rinkevich & Loya, 1979 and references therein |
| *Acanthaster* | *planci* | Echinodermata | Asteroidea | 21.75 | Nishida & Lucas, 1988; Yasuda et al., 2009; Hoegh-Guldberg & Pearse, 1995; Moran et al., 1992 | September | Koh Tao island, Thailand | Scott et al., 2015 |
| *Aquilonastra* | *burtoni* | Echinodermata | Asteroidea | 5 | James, 1972 | January-March | Marsa Mukebla, Elat, Red Sea | Achituv, 1973 |
| *Aquilonastra* | *coronata* | Echinodermata | Asteroidea | 20 | Komatsu, 1975 |  |  |  |
| *Archaster* | *typicus* | Echinodermata | Asteroidea | 25.75 | Mortensen, 1931; Komatsu et al., 2001 | Late June-July^1^; other regions spring-summer spawning is observed^2^; Mating season from September-October^3^ | Penghu, Taiwan^1^; Davao Gulf, Philippines^3^ | Run et al., 1988^1^; Keesing et al., 2011^2^; Bos et al., 2013^3^ |
| *Astropecten* | *polyacanthus* | Echinodermata | Asteroidea | 3.5 | Mortensen, 1937 |  |  |  |
| *Echinaster* | *purpureus* | Echinodermata | Asteroidea | 4 | Mortensen, 1938 |  |  |  |
| *Gomophia* | *egyptiaca* | Echinodermata | Asteroidea | 9 | Yamaguchi, 1974 |  |  |  |
| *Linckia* | *laevigata* | Echinodermata | Asteroidea | 22 | Yamaguchi, 1973 | Peak breeding in summer months in May-August^1^; near the equator may spawn throughout year^2^ | Guam, Micronesia^1^; multiple sites^2^ | Yamaguchi, 1977^1^; Pearse, 1968^2^ |
| *Luidia* | *maculata* | Echinodermata | Asteroidea | 64 | Komatsu et al., 1994 |  |  |  |
| *Parvulastra* | *amphitrite* | Echinodermata | Asteroidea | 0 | Hunt, 1993 |  |  |  |
| *Diadema* | *setosum* | Echinodermata | Echinoidea | 45 | Mortensen, 1937 | Probably spawn throught year at latitudes near equator, might be related to temperature^1^; August-December^2^ | multiple sites^1^; Gulf of Aqaba, Red Sea^2^ | Pearse, 1968^1^; Bronstein et al., 2016^2^ |
| *Echinometra* | *mathaei* | Echinodermata | Echinoidea | 40 | Onoda, 1936 | Probably spawn throught year at latitudes near equator^1^; Spawning peak from March-April^2^ | multiple sites^1^; Kenya^2^ | Pearse, 1968^1^; Muthiga et al., 2008^2^ |
| *Eucidaris* | *metularia* | Echinodermata | Echinoidea | 30 | Mortensen, 1937 |  |  |  |
| *Heterocentrotus* | *mammilatus* | Echinodermata | Echinoidea | 8 | Selkoe et al., 2014 |  |  |  |
| *Heterocentrotus* | *mammillatus* | Echinodermata | Echinoidea | 8 | Mortensen, 1937 |  |  |  |
| *Jackonaster* | *depressum* | Echinodermata | Echinoidea | 14 | Mortensen, 1938 |  |  |  |
| *Mespilia* | *globulus* | Echinodermata | Echinoidea | 30 | Onada, 1936; Emlet, 1995 |  |  |  |
| *Phyllacanthus* | *imperialis* | Echinodermata | Echinoidea | 3.5 | Olson et al., 1993 |  |  |  |
| *Prionocidaris* | *baculosa* | Echinodermata | Echinoidea | 25 | Mortensen, 1938 |  |  |  |
| *Salmacis* | *bicolor* | Echinodermata | Echinoidea | 23.5 | Aiyar, 1935 | Breeding throughout the year^1^; Congener *Salmacis virgulata* shows a spawning peak in the north-east monsoon from October-January^2^ | Madras^1^; Gulf of Mannar^2^ | Aiyar, 1935^1^; Saravanan et al., 2017^2^ |
| *Stomopneustes* | *variolaris* | Echinodermata | Echinoidea | 28 | Emlet, 2009 | Spawning may happen throughout the year, possible peak in March-April^1^; September-April^2^; December-February^3^ | Madras harbour^1^; South-west coast, India^2^; Oslo beach, South Africa^3^ | Giese et al., 1964^1^; Pillay, 1971^2^; Drummond, 1993^3^ |
| *Tripneustes* | *gratilla* | Echinodermata | Echinoidea | 18 | Mortensen, 1937 | Winter, early summer – April-October | Toliara, Madagascar | Vaïtilingon et al., 2005 |
| *Tripneustes spp.* |  | Echinodermata | Echinoidea | 18 | Lessios et al., 2003 |  |  |  |
| *Actinopyga* | *echinites* | Echinodermata | Holothuroidea | 18 | Chen & Chian, 1990 | Major spawning events in December-January, minor event in April-May | La Réunion | Kohler et al., 2010 |
| *Holothuria* | *nobilis* | Echinodermata | Holothuroidea | 28 | Ulthicke & Benzie, 2000 | July-September | Minicoy, Lakshadweep islands | Kandan, 1994 |
| *Holothuria* | *spinifera* | Echinodermata | Holothuroidea | 12.5 | Asha, 2005; Asha & Muthiah, 2008 | November-March based on gonadal observations | Individuals obtained from Koswari and Van Island, Tuticorin | Asha & Muthiah, 2008 |
| *Holothuria* | *atra* | Echinodermata | Holothuroidea | 15 | Selkoe et al., 2014 | July-August, February-March | Tuticorin | Ram Mohan, 2001 |
| *Holothuria* | *arenicola* | Echinodermata | Holothuroidea | 28 | Mortensen, 1938 |  |  |  |
| *Holothuria* | *difficilis* | Echinodermata | Holothuroidea | 12 | Mortensen, 1938 | August-September | Nanwan, Wanlitung, southern Taiwan | Chao et al., 1995 |
| *Holothuria* | *impatiens* | Echinodermata | Holothuroidea | 14 | Mortensen, 1938 | Congener *Holothuria leucospilota* spawns from October-January, June-September | Anjuna, Goa | Jayasree & Bhavanarayana, 1994 |
| *Holothuria* | *scabra* | Echinodermata | Holothuroidea | 15.33 | James, 2004; Ramofafia Ramofafia, Byrne & Battaglene, 2003 | Throughout the year, peaks from March-April, July, September-October | India | James, 1989 |
| *Breviturma* | *pica* | Echinodermata | Ophiuroidea | 50 | Selkoe et al., 2014 |  |  |  |
| *Ophiocoma* | *erinaceus* | Echinodermata | Ophiuroidea | 50 | Selkoe et al., 2014 | Throughout the year in *Ophiocoma scolopendrina*, *O. venosa* from February-May | Toliara, Madagascar | Delroisse et al., 2013 |
| *Ophiomaza* | *cacaotica* | Echinodermata | Ophiuroidea | 13 | Mortensen, 1938 |  |  |  |
| *Gafrarium* | *pectinatum* | Mollusca | Bivalvia | 13 | Jagadis, 2011 | For congener *Gafrarium tumidum*, continuous breeding but peak spawning in November and minor spawning in April | Chinnapalam, Pamban | Jagadis & Rajagopal, 2007 |
| *Magallana* | *bilineata* | Mollusca | Bivalvia | 19 | Nayar et al., 1982; Rao, 1983 | Breeding season from November-December | Individuals obtained from Adyar and Ennore estuaries, Chennai | Rao, 1983 |
| *Meretrix* | *casta* | Mollusca | Bivalvia | 8.5 | Sreenivasan & Rao, 1991 | Breeding peaks in summer months, September-November; Mandapam has summer break in spawning^1^; Spawning peaks in April-September^2^; May, September-October^3^ | Beypore, Kozhapura^1^; Vellar estuary^2^; Muttukadu^3^ | Seshappa, 1967 and references therein^1^; references within Jagadis & Rajagopal, 2007^2,3^ |
| *Meretrix* | *meretrix* | Mollusca | Bivalvia | 8.5 | Narasimham et al., 1988 | Breeding for upto 9 months, absence of spawning from November-December |  | Narasimham et al., 1988 |
| *Perna* | *viridis* | Mollusca | Bivalvia | 17.5 | Rao, Kumari & Qasim, 1976; Benson et al., 2001 ; Barber et al., 2005 | Reported to possibly vary geographically. On east coast, before the summer monsoon (October-July) and on the west coast from July-December |  | References within Nair, 2001 |
| *Perna* | *indica* | Mollusca | Bivalvia |  |  | May-September with a peak from July-August | Vizhinjam | Appukuttan & Nair, 1983 |
| *Pinctada* | *fucata* | Mollusca | Bivalvia | 28.5 | Alagarswami et al., 1983 | Peaks in spontaneous spawning from November-January, July-August | Individuals obtained from pearl culture farms, Kochi | Alagarswami et al., 1983 |
| *Placuna* | *placenta* | Mollusca | Bivalvia |  |  | February-May and October-December with some variation between years | Individuals obtained from Kakinada Bay | Narasimham, 1984 |
| *Protapes* | *gallus* | Mollusca | Bivalvia | 10.5 | Gireesh & Gopinathan, 2004 |  |  |  |
| *Saccostrea* | *cuccullata* | Mollusca | Bivalvia | 20 | Sukumar and Joseph, 1988 | Continuous breeding season, peaks in December-January, minor peak in May-June. Studies in Someshwar coast indicate prolonged soawning from June-December, in Shirgaon creek from November-December. | Ashtamudi lake, Kerala | Kripa & Salih, 1996 and references therein |
| *Tegillarca* | *granosa* | Mollusca | Bivalvia | 17 | Muthiah et al., 1992 | Spawns throughout the year with peaks in January and April^1^; Peak spawning varies with location and year but between February-March, May-June and October-November^2^ | Kakinada Bay^1,2^ | Narasimham, 1988 and references therein |
| *Tridacna* | *maxima* | Mollusca | Bivalvia | 9 | Benzie, 1994; Benzie & Williams, 1997 | For *Tridacna* in general, spawning may be related to latitude, where seasonality may be observed at higher latitudes |  | Lucas, 1994; Soo & Todd, 2014 |
| *Tridacna* | *crocea* | Mollusca | Bivalvia | 10.13 | Kochzius & Nuryanto, 2008; DeBoer et al., 2008; Benzie & Williams, 1997 | For *Tridacna* in general, spawning may be related to latitude, where seasonality may be observed at higher latitudes |  | Lucas, 1994; Soo & Todd, 2014 |
| *Babylonia* | *spirata* | Mollusca | Gastropoda | 17 | Sreejaya et al., 2004 | January-April with peaks in January and April, September-December | Individuals obtained from trawl landing, Neendakara | Sreejaya et al., 2004 |
| *Chicoreus* | *ramosus* | Mollusca | Gastropoda | 27 | Jagadis et al, 2017 | January-March^1^;  November-March^2^ | Anjuna, Goa^1^; Mandapam^2^ | Jagadis et al., 2017^1^; Ramesh et al., 1992^2^ |
| *Conus* | *biliosus* | Mollusca | Gastropoda | 26 | Zehra & Perveen, 1991 | *Conus biliosus* from April-May and congener *C. coronatus* spawns from April-June with peak in April^1^; Congener *C. araneosus* observed to lay egg capsules from October-March^2^ | Buleji, Karachi^1^; Gulf of Mannar and Palk Bay^2^ | Zehra & Perveen, 1991^1^; Natarajan, 1957^2^ |
| *Cypraea* | *tigris* | Mollusca | Gastropoda |  |  | October (but spawning only observed once) | Landing centre, Tuticorin | Jagadis et al, 2017 |
| *Drupella* | *cornus* | Mollusca | Gastropoda | 21 | Johnson et al., 1993 |  |  |  |
| *Echinolittorina* |  | Mollusca | Gastropoda | 24.5 | Reid, 2007 | March-June for *Echinolittorina trochoides* | Penang coast, Malaysia | Berry, 1986 |
| *Lambis* | *lambis* | Mollusca | Gastropoda | 16 | Jagadis et al., 2012 | October-December | Individuals obtained from landing centres in Vellapatti and Sippikulam, Tuticorin | Jagadis et al., 2012 |
| *Littoraria* |  | Mollusca | Gastropoda | 45.5 | Reid et al, 2010 | Throughout the year in *Littoraria scabra* | Porto Novo, Cuddalore | Marimuthu & Kasinathan, 1986 |
| *Pirenella* | *cingulata* | Mollusca | Gastropoda | 12.5 | Sreenivasan, 1985 | January-September | Vellar estuary | Sreenivasan, 1985 |

<http://eprints.cmfri.org.in/id/eprint/449>
