## Supporting Information Appendix S2 for "Connectivity networks and delineation of distinct coastal provinces along the Indian coastline using large-scale Lagrangian transport simulations"

**Appendix 2.** Description of particle release locations and settlement polygons used to evaluate coastal connectivity.

**Particle release locations**

Interpolation of current velocities in the general circulation model output can lead to lower than expected velocity values in grid cells immediately adjacent to the land mask. To prevent erroneous particle retention in these cells because of low velocities, we moved the release locations to the second cell from the land mask, at a distance of one and a half cell-widths (1/8°) from the HYCOM land mask into the sea. Our simulations, therefore, model the trajectories of particles that have moved away from the coast and do not effectively model self-recruitment and retention at the coast due to small-scale oceanographic processes such as tidal flows.

**Settlement polygons**

Successful dispersal was recorded when particles reached settlement polygons along the coast at the end of their pelagic larval duration (PLD). The settlement polygons were placed one cell into the sea, away from the HYCOM land-mask. This was done to avoid considering particles that may be retained close to the coast because of low velocities at cells immediately adjacent to the land-mask as explained above. Most (500 out of 528) of the settlement polygons encompassed four release locations and had a mean area of 191.53 km^2^ (range: 45.70-321.03 km^2^) (Figure S2.3). To account for the variation in polygon area available for settlement across coastal polygons, the transfer probability was calculated per unit surface area.

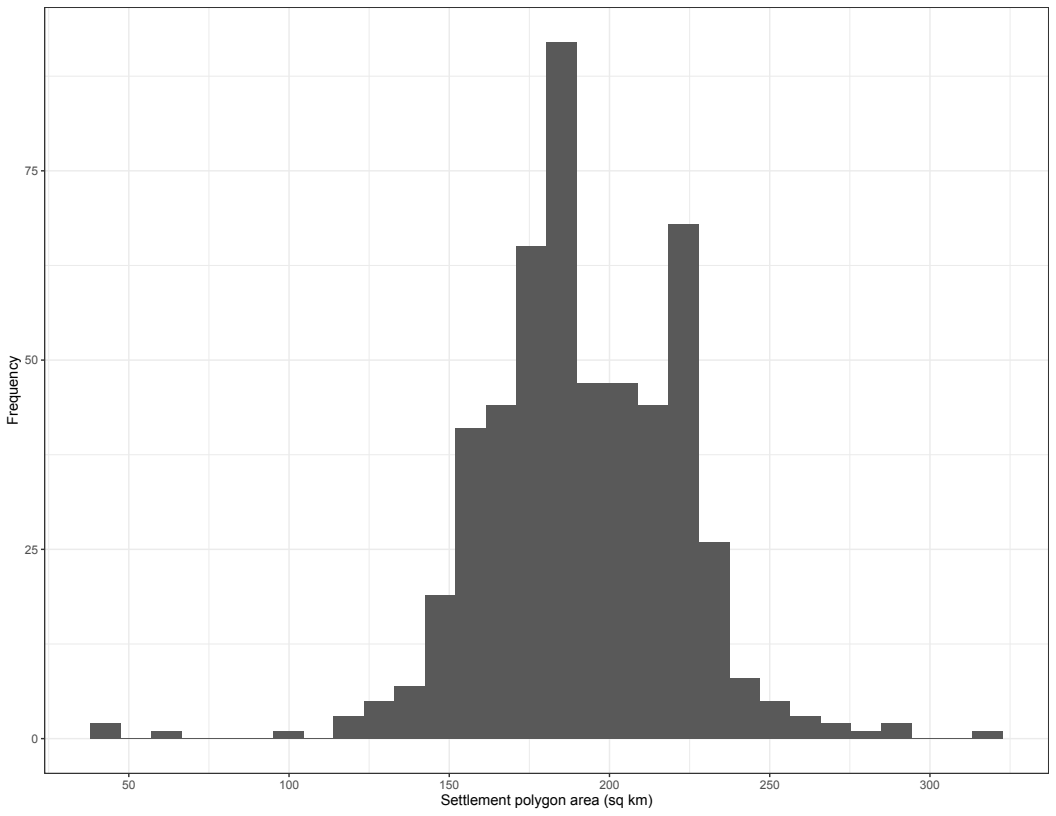

**Figure S2.3.** Frequency distribution of settlement polygon area used in the current study.
