## Supporting Information Appendix S3 for "Connectivity networks and delineation of distinct coastal provinces along the Indian coastline using large-scale Lagrangian transport simulations"

**Appendix 3.** Classification of pelagic larval duration (PLD) classes for connectivity analysis.

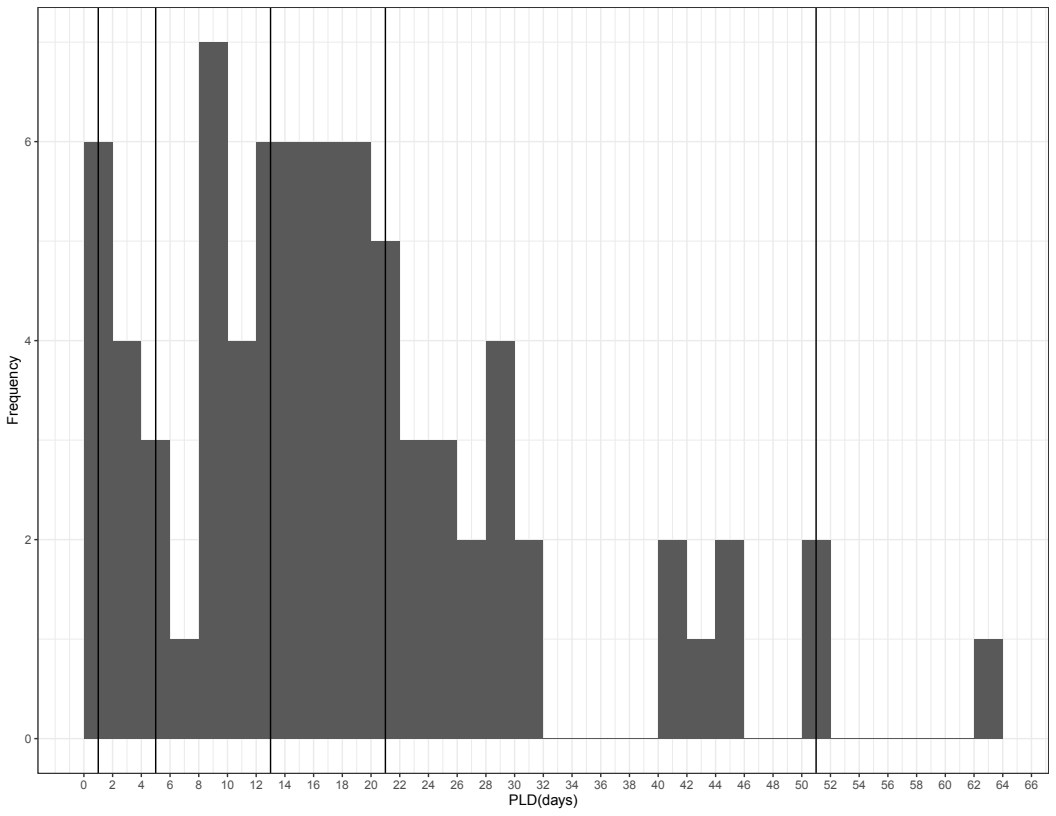

**Figure S3.4.** Distribution of PLD across marine invertebrates found in the coastal Indian Ocean, with vertical lines representing the range of PLD values within each PLD-class that was used for connectivity analysis (2-4 days, 6-12 days, 14-20 days and 22-50 days). A PLD range of 2-50 days with two-day interval was used for this analysis. Note that this histogram omits one record of PLD 240 days from *Panulirus homarus.*

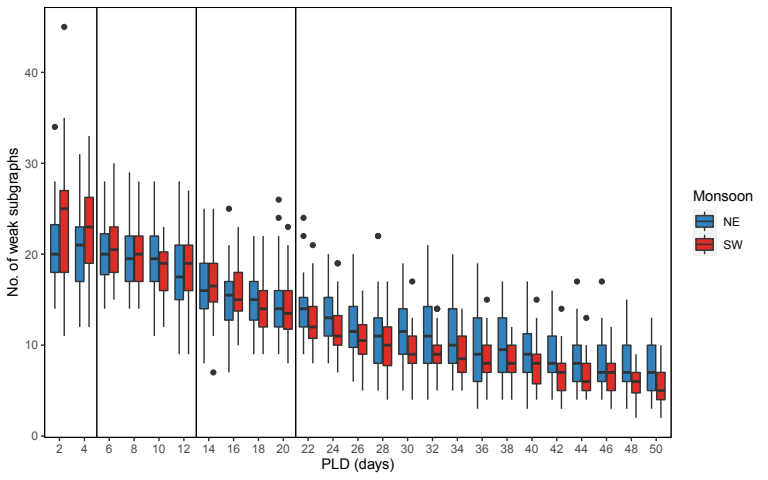

**Figure S3.5.** Distribution of weak subgraphs by PLD for different seasons. The vertical lines represent the range of PLD values within each PLD-class that was used for connectivity analysis.
