## Supporting Information Appendix S4 for "Connectivity networks and delineation of distinct coastal provinces along the Indian coastline using large-scale Lagrangian transport simulations"

**Appendix 4.** Statistical tests to evaluate differences in the distribution of proportion connectance, number of singletons, weak subgraphs, strong subgraphs, Infomap communities and coherence ratio (rho) values with season and pelagic larval duration (PLD) class.

**Table S4.2.** Pairwise Kolmogorov-Smirnov test to compare differences in the cumulative frequency distribution of proportion connectance with season and PLD-class. A Bonferroni correction was applied to the P-value (referred to in the table as ‘corrected’) to account for multiple comparisons.

| **Comparison** | **Test statistic (D)** | **P-value** | **P-value (corrected)** | **Significance** |
| --- | --- | --- | --- | --- |
| **Year** | | | | |
| 2009 vs 2010 | 0.1667 | 0.1389 | 0.4168 |  |
| 2009 vs 2011 | 0.1042 | 0.6749 | 1.0000 |  |
| 2010 vs 2011 | 0.1667 | 0.1389 | 0.4168 |  |
| **Season** | | | | |
| ne vs sw | 0.3264 | 4.35 × 10-07 | 4.35 × 10-07 | *** |
| **PLD:Season** | | | | |
| ne_2-4 vs ne_6-12 | 0.7778 | 6.97 × 10-10 | 1.95 × 10-08 | *** |
| ne_2-4 vs ne_14-20 | 0.7222 | 1.40 × 10-08 | 3.92 × 10-07 | *** |
| ne_2-4 vs ne_22-50 | 1.0000 | 4.44 × 10-16 | 1.24 × 10-14 | *** |
| ne_6-12 vs ne_14-20 | 0.1389 | 8.78 × 10-01 | 1.00 × 10+00 |  |
| ne_6-12 vs ne_22-50 | 0.9722 | 3.33 × 10-15 | 9.33 × 10-14 | *** |
| ne_14-20 vs ne_22-50 | 0.9722 | 3.33 × 10-15 | 9.33 × 10-14 | *** |
| sw_2-4 vs sw_6-12 | 0.3333 | 3.66 × 10-02 | 1.00 × 10+00 |  |
| sw_2-4 vs sw_14-20 | 0.2222 | 3.36 × 10-01 | 1.00 × 10+00 |  |
| sw_2-4 vs sw_22-50 | 1.0000 | 0.0000 | 1.24 × 10-14 | *** |
| sw_6-12 vs sw_14-20 | 0.1667 | 6.99 × 10-01 | 1.00 × 10+00 |  |
| sw_6-12 vs sw_22-50 | 1.0000 | 4.44 × 10-16 | 1.24 × 10-14 | *** |
| sw_14-20 vs sw_22-50 | 0.9722 | 3.33 × 10-15 | 9.33 × 10-14 | *** |
| ne_2-4 vs sw_2-4 | 0.4722 | 6.53 × 10-04 | 1.83 × 10-02 | . |
| ne_6-12 vs sw_6-12 | 0.5833 | 9.57 × 10-06 | 2.68 × 10-04 | ** |
| ne_14-20 vs sw_14-20 | 0.6389 | 8.30 × 10-07 | 2.33 × 10-05 | *** |
| ne_22-50 vs sw_22-50 | 0.3333 | 3.66 × 10-02 | 1.00 × 10+00 |  |

**Table S4.3.** Pairwise Kolmogorov-Smirnov test to compare differences in the cumulative frequency distribution of number of singletons with season and PLD-class. A Bonferroni correction was applied to the P-value (referred to in the table as ‘corrected’) to account for multiple comparisons.

| **Comparison** | **Test statistic (D)** | **P-value** | **P-value (corrected)** | **Significance** |
| --- | --- | --- | --- | --- |
| **Year** | | | | |
| 2009 vs 2010 | 0.1600 | 4.27 × 10-07 | 1.28 × 10-06 | *** |
| 2009 vs 2011 | 0.0617 | 2.04 × 10-01 | 6.12 × 10-01 |  |
| 2010 vs 2011 | 0.1850 | 2.41 × 10-09 | 7.24 × 10-09 | *** |
| **Season** | | | | |
| ne vs sw | 0.3544 | 0 | 0 | *** |
| **PLD:Season** | | | | |
| ne_2-4 vs ne_6-12 | 0.6111 | 2.90 × 10-06 | 8.12 × 10-05 | *** |
| ne_2-4 vs ne_14-20 | 0.8333 | 2.78 × 10-11 | 7.78 × 10-10 | *** |
| ne_2-4 vs ne_22-50 | 0.4167 | 3.86 × 10-03 | 1.08 × 10-01 |  |
| ne_6-12 vs ne_14-20 | 0.5833 | 9.57 × 10-06 | 2.68 × 10-04 | ** |
| ne_6-12 vs ne_22-50 | 0.2778 | 1.24 × 10-01 | 1.00 × 10+00 |  |
| ne_14-20 vs ne_22-50 | 0.4722 | 6.53 × 10-04 | 1.83 × 10-02 | . |
| sw_2-4 vs sw_6-12 | 0.5833 | 9.57 × 10-06 | 2.68 × 10-04 | ** |
| sw_2-4 vs sw_14-20 | 0.8333 | 2.78 × 10-11 | 7.78 × 10-10 | *** |
| sw_2-4 vs sw_22-50 | 0.2778 | 1.24 × 10-01 | 1.00 × 10+00 |  |
| sw_6-12 vs sw_14-20 | 0.4167 | 3.86 × 10-03 | 1.08 × 10-01 |  |
| sw_6-12 vs sw_22-50 | 0.4722 | 6.53 × 10-04 | 1.83 × 10-02 | . |
| sw_14-20 vs sw_22-50 | 0.7500 | 3.21 × 10-09 | 8.99 × 10-08 | *** |
| ne_2-4 vs sw_2-4 | 0.4167 | 3.86 × 10-03 | 1.08 × 10-01 |  |
| ne_6-12 vs sw_6-12 | 0.6667 | 2.25 × 10-07 | 6.30 × 10-06 | *** |
| ne_14-20 vs sw_14-20 | 0.4167 | 3.86 × 10-03 | 1.08 × 10-01 |  |
| ne_22-50 vs sw_22-50 | 0.3056 | 6.94 × 10-02 | 1.00 × 10+00 |  |

**Table S4.4.** Pairwise Kolmogorov-Smirnov test to compare differences in the cumulative frequency distribution of number of weak subgraphs with season and PLD-class. A Bonferroni correction was applied to the P-value (referred to in the table as ‘corrected’) to account for multiple comparisons.

| **Comparison** | **Test statistic (D)** | **P-value** | **P-value (corrected)** | **Significance** |
| --- | --- | --- | --- | --- |
| **Year** | | | | |
| 2009 vs 2010 | 0.1146 | 0.5542 | 1 |  |
| 2009 vs 2011 | 0.0833 | 0.8928 | 1 |  |
| 2010 vs 2011 | 0.1042 | 0.6749 | 1 |  |
| **Season** | | | | |
| ne vs sw | 0.1111 | 0.3364 | 0.3364 |  |
| **PLD:Season** | | | | |
| ne_2-4 vs ne_6-12 | 0.2500 | 2.11 × 10-01 | 1.00 × 10+00 |  |
| ne_2-4 vs ne_14-20 | 0.3056 | 6.94 × 10-02 | 1.00 × 10+00 |  |
| ne_2-4 vs ne_22-50 | 0.9722 | 3.33 × 10-15 | 9.33 × 10-14 | *** |
| ne_6-12 vs ne_14-20 | 0.1111 | 9.79 × 10-01 | 1.00 × 10+00 |  |
| ne_6-12 vs ne_22-50 | 0.9167 | 1.46 × 10-13 | 4.08 × 10-12 | *** |
| ne_14-20 vs ne_22-50 | 0.9167 | 1.46 × 10-13 | 4.08 × 10-12 | *** |
| sw_2-4 vs sw_6-12 | 0.3611 | 1.83 × 10-02 | 5.12 × 10-01 |  |
| sw_2-4 vs sw_14-20 | 0.5833 | 9.57 × 10-06 | 2.68 × 10-04 | ** |
| sw_2-4 vs sw_22-50 | 1.0000 | 4.44 × 10-16 | 1.24 × 10-14 | *** |
| sw_6-12 vs sw_14-20 | 0.3333 | 3.66 × 10-02 | 1.00 × 10+00 |  |
| sw_6-12 vs sw_22-50 | 1.0000 | 4.44 × 10-16 | 1.24 × 10-14 | *** |
| sw_14-20 vs sw_22-50 | 1.0000 | 4.44 × 10-16 | 1.24 × 10-14 | *** |
| ne_2-4 vs sw_2-4 | 0.1944 | 5.04 × 10-01 | 1.00 × 10+00 |  |
| ne_6-12 vs sw_6-12 | 0.1389 | 8.78 × 10-01 | 1.00 × 10+00 |  |
| ne_14-20 vs sw_14-20 | 0.2500 | 2.11 × 10-01 | 1.00 × 10+00 |  |
| ne_22-50 vs sw_22-50 | 0.4444 | 1.63 × 10-03 | 4.57 × 10-02 | . |

**Table S4.5.** Pairwise Kolmogorov-Smirnov test to compare differences in the cumulative frequency distribution of number of strong subgraphs with season and PLD-class. A Bonferroni correction was applied to the P-value (referred to in the table as ‘corrected’) to account for multiple comparisons.

| **Comparison** | **Test statistic (D)** | **P-value** | **P-value (corrected)** | **Significance** |
| --- | --- | --- | --- | --- |
| **Year** | | | | |
| 2009 vs 2010 | 0.1250 | 0.4413 | 1.0000 |  |
| 2009 vs 2011 | 0.0729 | 0.9605 | 1.0000 |  |
| 2010 vs 2011 | 0.1979 | 0.0465 | 0.1396 |  |
| **Season** | | | | |
| ne vs sw | 0.25 | 0.0003 | 0.0002 | ** |
| **PLD:Season** | | | | |
| ne_2-4 vs ne_6-12 | 0.7222 | 1.40 × 10-08 | 3.92 × 10-07 | *** |
| ne_2-4 vs ne_14-20 | 0.9722 | 3.33 × 10-15 | 9.33 × 10-14 | *** |
| ne_2-4 vs ne_22-50 | 1.0000 | 4.44 × 10-16 | 1.24 × 10-14 | *** |
| ne_6-12 vs ne_14-20 | 0.6944 | 5.77 × 10-08 | 1.62 × 10-06 | *** |
| ne_6-12 vs ne_22-50 | 0.9444 | 2.26 × 10-14 | 6.34 × 10-13 | *** |
| ne_14-20 vs ne_22-50 | 0.6944 | 5.77 × 10-08 | 1.62 × 10-06 | *** |
| sw_2-4 vs sw_6-12 | 0.6944 | 5.77 × 10-08 | 1.62 × 10-06 | *** |
| sw_2-4 vs sw_14-20 | 0.8889 | 8.87 × 10-13 | 2.48 × 10-11 | *** |
| sw_2-4 vs sw_22-50 | 0.9722 | 3.33 × 10-15 | 9.33 × 10-14 | *** |
| sw_6-12 vs sw_14-20 | 0.5000 | 2.47 × 10-04 | 6.91 × 10-03 | * |
| sw_6-12 vs sw_22-50 | 0.7778 | 6.97 × 10-10 | 1.95 × 10-08 | *** |
| sw_14-20 vs sw_22-50 | 0.5278 | 8.83 × 10-05 | 2.47 × 10-03 | * |
| ne_2-4 vs sw_2-4 | 0.3889 | 8.64 × 10-03 | 2.42 × 10-01 |  |
| ne_6-12 vs sw_6-12 | 0.5278 | 8.83 × 10-05 | 2.47 × 10-03 | * |
| ne_14-20 vs sw_14-20 | 0.6111 | 2.90 × 10-06 | 8.12 × 10-05 | *** |
| ne_22-50 vs sw_22-50 | 0.3611 | 1.83 × 10-02 | 5.12 × 10-01 |  |

**Table S4.6.** Pairwise Kolmogorov-Smirnov test to compare differences in the cumulative frequency distribution of number of Infomap communities (>1 membership) with season and PLD-class. A Bonferroni correction was applied to the P-value (referred to in the table as ‘corrected’) to account for multiple comparisons.

| **Comparison** | **Test statistic (D)** | **P-value** | **P-value (corrected)** | **Significance** |
| --- | --- | --- | --- | --- |
| **Year** | | | | |
| 2009 vs 2010 | 0.0938 | 0.7928 | 1 |  |
| 2009 vs 2011 | 0.0729 | 0.9605 | 1 |  |
| 2010 vs 2011 | 0.0833 | 0.8928 | 1 |  |
| **Season** | | | | |
| ne vs sw | 0.1944 | 0.0086 | 0.0086 | * |
| **PLD:Season** | | | | |
| ne_2-4 vs ne_6-12 | 0.9722 | 3.33 × 10-15 | 9.33 × 10-14 | *** |
| ne_2-4 vs ne_14-20 | 1.0000 | 4.44 × 10-16 | 1.24 × 10-14 | *** |
| ne_2-4 vs ne_22-50 | 1.0000 | 4.44 × 10-16 | 1.24 × 10-14 | *** |
| ne_6-12 vs ne_14-20 | 0.6389 | 8.30 × 10-07 | 2.33 × 10-05 | *** |
| ne_6-12 vs ne_22-50 | 1.0000 | 4.44 × 10-16 | 1.24 × 10-14 | *** |
| ne_14-20 vs ne_22-50 | 0.8611 | 5.10 × 10-12 | 1.43 × 10-10 | *** |
| sw_2-4 vs sw_6-12 | 1.0000 | 4.44 × 10-16 | 1.24 × 10-14 | *** |
| sw_2-4 vs sw_14-20 | 1.0000 | 4.44 × 10-16 | 1.24 × 10-14 | *** |
| sw_2-4 vs sw_22-50 | 1.0000 | 4.44 × 10-16 | 1.24 × 10-14 | *** |
| sw_6-12 vs sw_14-20 | 0.5278 | 8.83 × 10-05 | 2.47 × 10-03 | * |
| sw_6-12 vs sw_22-50 | 0.9722 | 3.33 × 10-15 | 9.33 × 10-14 | *** |
| sw_14-20 vs sw_22-50 | 0.8889 | 8.87 × 10-13 | 2.48 × 10-11 | *** |
| ne_2-4 vs sw_2-4 | 0.4167 | 3.86 × 10-03 | 1.08 × 10-01 |  |
| ne_6-12 vs sw_6-12 | 0.5000 | 2.47 × 10-04 | 6.91 × 10-03 | * |
| ne_14-20 vs sw_14-20 | 0.3611 | 1.83 × 10-02 | 5.12 × 10-01 |  |
| ne_22-50 vs sw_22-50 | 0.4167 | 3.86 × 10-03 | 1.08 × 10-01 |  |

**Table S4.7.** Pairwise Kolmogorov-Smirnov test to compare differences in the cumulative frequency distribution of coherence ratio (rho) values of Infomap communities with season and PLD-class. A Bonferroni correction was applied to the P-value (referred to in the table as ‘corrected’) to account for multiple comparisons.

| **Comparison** | **Test statistic (D)** | **P-value** | **P-value (corrected)** | **Significance** |
| --- | --- | --- | --- | --- |
| **Year** | | | | |
| 2009 vs 2010 | 0.01844 | 0.4805 | 1 |  |
| 2009 vs 2011 | 0.01973 | 0.3987 | 1 |  |
| 2010 vs 2011 | 0.01459 | 0.7653 | 1 |  |
| **Season** | | | | |
| ne vs sw | 0.0608 | 2.16 × 10-10 | 2.16 × 10-10 | *** |
| **PLD:Season** | | | | |
| ne_2_4 vs ne_6_12 | 0.0878 | 0.00 | 8.08 × 10-06 | *** |
| ne_2_4 vs ne_14_20 | 0.1276 | 0.00 | 1.41 × 10-11 | *** |
| ne_2_4 vs ne_22_50 | 0.2211 | 0.00 | 0.00 × 10+00 | *** |
| ne_6_12 vs ne_14_20 | 0.0565 | 0.01 | 3.69 × 10-01 |  |
| ne_6_12 vs ne_22_50 | 0.1570 | 0.00 | 1.71 × 10-12 | *** |
| ne_14_20 vs ne_22_50 | 0.1326 | 0.00 | 8.25 × 10-08 | *** |
| sw_2_4 vs sw_6_12 | 0.0574 | 0.01 | 1.47 × 10-01 |  |
| sw_2_4 vs sw_14_20 | 0.1394 | 0.00 | 1.42 × 10-12 | *** |
| sw_2_4 vs sw_22_50 | 0.3966 | 0.00 | 0.00 × 10+00 | *** |
| sw_6_12 vs sw_14_20 | 0.0915 | 0.00 | 5.79 × 10-04 | ** |
| sw_6_12 vs sw_22_50 | 0.3467 | 0.00 | 0.00 × 10+00 | *** |
| sw_14_20 vs sw_22_50 | 0.2741 | 0.00 | 0.00 × 10+00 | *** |
| ne_2_4 vs sw_2_4 | 0.0627 | 0.00 | 5.98 × 10-03 | * |
| ne_6_12 vs sw_6_12 | 0.0405 | 0.14 | 1.00 × 10+00 |  |
| ne_14_20 vs sw_14_20 | 0.0589 | 0.02 | 5.54 × 10-01 |  |
| ne_22_50 vs sw_22_50 | 0.2221 | 0.00 | 0.00 × 10+00 | *** |
